# Tissue-specific modulation of NADH consumption as an anti-aging intervention in Drosophila

**DOI:** 10.1101/2025.01.06.631511

**Authors:** Shweta Yadav, Xingxiu Pan, Shengxi Li, Paige LaRae Martin, Ngoc Hoang, Kejin Chen, Aditi Karhadkar, Jatin Malhotra, Austin L. Zuckerman, Subrata Munan, Markus K. Klose, Lin Wang, Valentin Cracan, Andrey A Parkhitko

## Abstract

Aging is characterized by extensive metabolic dysregulation. Redox coenzyme nicotinamide adenine dinucleotide (NAD) can exist in oxidized (NAD^+^) or reduced (NADH) states, which together form a key NADH/NAD^+^ redox pair. Total levels of NAD decline with age in a tissue-specific manner, thereby playing a significant role in the aging process. Supplementation with NAD precursors boosts total cellular NAD levels and provides some therapeutic benefits in human clinical trials. However, supplementation studies cannot determine tissue-specific effects of an altered NADH/NAD^+^ ratio. Here, we created transgenic *Drosophila* expressing a genetically encoded xenotopic tool *Lb*NOX to directly manipulate the cellular NADH/NAD^+^ ratio. We found that *Lb*NOX expression in *Drosophila* impacts both NAD(H) and NADP(H) metabolites in a sex-specific manner. *Lb*NOX rescues neuronal cell death induced by the expression of mutated alpha-B crystallin in the *Drosophila* eye, a widely used system to study reductive stress. Utilizing *Lb*NOX, we demonstrate that targeting redox NAD metabolism in different tissues may have drastically different outcomes, as the expression of *Lb*NOX solely in the muscle is much more effective for rescuing paraquat-induced oxidative stress compared to whole-body expression. Excitingly, we demonstrate that perturbing NAD(P) metabolism in non-neuronal tissues is sufficient to rejuvenate sleep profiles in aged flies to a youthful state. In summary, we used xenotopic tool *Lb*NOX to identify tissues and metabolic processes which benefited the most from the modulation of the NAD metabolism thereby highlighting important aspects of rebalancing the NAD and NADP pools, all of which can be translated into novel designs of NAD-related human clinical trials.

**Significance statement:** Total levels of NAD decline with age in a tissue-specific manner, thereby playing a significant role in the aging process. Supplementation with NAD precursors boosts organismal NAD levels but cannot determine tissue-specific effects of altered NAD metabolism. Here, we created transgenic *Drosophila* expressing a genetically encoded xenotopic tool, *Lb*NOX, to directly manipulate NAD metabolism. We demonstrate that targeting NAD metabolism in just one tissue may be more effective than altering whole-body metabolism and can reverse some aging-related manifestations in a sex-specific manner. We anticipate that our work will define the tissues that benefit the most from targeting NAD metabolism and aid in designing better human clinical trials.

## Introduction

Metabolic dysregulation represents one of the major driving forces in aging, leading to impaired organismal fitness, an age-dependent increase in susceptibility to diseases, a decreased ability to mount a stress response, and increased frailty (1). Dysregulation of nicotinamide adenine dinucleotide (NAD) metabolism has emerged as a contributing factor in the pathogenesis of aging and multiple age-related diseases (2–4). NAD in cells is found in both oxidized (NAD^+^) and reduced (NADH) forms, which together furnish cells with a key NADH/NAD^+^ redox pair that sits at the core of redox metabolism and signaling (5). In addition, NAD^+^ also serves as an essential substrate for non-redox NAD^+^-dependent enzymes, including sirtuins, CD38, ARTs, SARM1, and poly (ADP-ribose) polymerases (5, 6).

The total NAD pool declines with age and under various pathological conditions in a tissue-specific manner, potentially contributing to the exacerbation of the pathological state. However, whether this occurs universally across different species and tissues remains unclear (7). Multiple preclinical studies in rodents provide evidence of the beneficial effects of supplementation with NAD biosynthetic precursors that boost total cellular NAD levels (7–9). However, supplementation studies do not provide an assessment of the tissue-specific role of restoring NAD levels or how does the NADH/NAD^+^ redox potential itself regulates the aging process.

Here, we used an alternative approach to directly manipulate NADH/NAD^+^ ratio by introducing an enzyme from a different species (“xenotopic approach”). We employed *L. brevis* H_2_O-froming NADH oxidase (*Lb*NOX) (10) as a genetically encoded xenotopic tool previously used for a compartment-specific decrease of the NADH/NAD^+^ ratio in living cells. We reasoned that a pro-oxidative shift (a decrease) in NADH/NAD^+^ ratio would be beneficial as an elevated NADH/NAD^+^ ratio (sometimes referred to as NADH-reductive stress) is linked to various pathological states ranging from primary mitochondrial diseases to neurodegeneration, as well as to aging-associated metabolic changes (11, 12). There are existing genetic tools to express transgenes in *Drosophila,* which allows for targeted expression of *Lb*NOX in different tissues or population of cells. The application of this model system can directly address how manipulation of cellular NADH/NAD^+^ ratio affects various organismal processes.

We show that cytosolic *Lb*NOX expression in transgenic *Drosophila* impacts both NADH/NAD^+^ and NADPH/NADP^+^ ratios to a different extent and in a sex-dependent manner. We further show that *Lb*NOX expression promotes resistance to oxidative and starvation stress, extends lifespan, and that targeting NAD metabolism in different tissues may have drastically different outcomes. Finally, we demonstrate that tissue-specific *Lb*NOX expression rejuvenates sleep profiles in aged flies back to a youthful state.

In summary, we demonstrate the broad impacts of modulating NADH-consuming activity using the xenotopic tool *Lb*NOX in the context of multiple aging-associated metabolic changes. We anticipate that our work will further clarify tissue-specific roles of NAD metabolism in the aging process and aid in the design of “combinatorial” clinical trials that can target both redox-neutral and redox-dependent aspects of NAD metabolism or combine NAD precursors with xenotopic tools.

## Results

### *Lb*NOX expression alters NAD(P)H/NAD(P)^+^ ratios in *Drosophila* in a sex-dependent manner

NAD boosters/precursors supplementation can increase organismal NAD levels; however, supplementation studies do not allow testing of tissue-specific roles of an altered NADH/NAD^+^ ratio in regulating different biological processes and cannot distinguish the impact of redox reactions from non-redox pathways. Therefore, we turned our attention to a *L. brevis* derived H_2_O-forming NADH oxidase (*Lb*NOX) as a genetically encoded xenotopic tool for inducing a compartment-specific decrease in the NADH/NAD^+^ ratio (10). *Lb*NOX catalyzes the conversion of NADH to NAD^+^ generating water as a byproduct (**Figure 1A**) (10). We cloned *Lb*NOX into the pWALIUM10-ROE *Drosophila* expression vector and created transgenic flies carrying the *Lb*NOX transgene on the 2nd (line #128) or on the 3rd (line #129) chromosome at a defined genetic locus. To confirm the expression of *Lb*NOX in flies, we expressed *Lb*NOX in adults ubiquitously using the *tubulin-Gal4, tubulin-Gal80^ts^* temperature-inducible system (13). Gal80^ts^ is active at 18°C and represses Gal4; while at 29°C, Gal80^ts^ is inactivated, allowing for the Gal4-dependent expression of *Lb*NOX. Genetic crosses were set at 18°C and maintained until the progeny eclosed followed by switching to 29°C to induce expression of *Lb*NOX for 7 days. As a control, we used the same parental line that we had injected with the UAS-*Lb*NOX construct to produce the transgenic UAS-*Lb*NOX flies. We confirmed expression of *Lb*NOX in flies by Western blotting (**Figure 1B**). Although NAD metabolites are known to be compartmentalized within the cell, due to technical difficulties, we could not create transgenic flies with a mitochondria-targeted form of *Lb*NOX. As we were interested in assessing the effects of altering NADH/NAD^+^ ratio in aged flies and wanted to explore sex-specific differences, we confirmed that the expression level of *Lb*NOX is similar across male and female flies and at different ages (**Figure 1B**).

**Figure 1.**
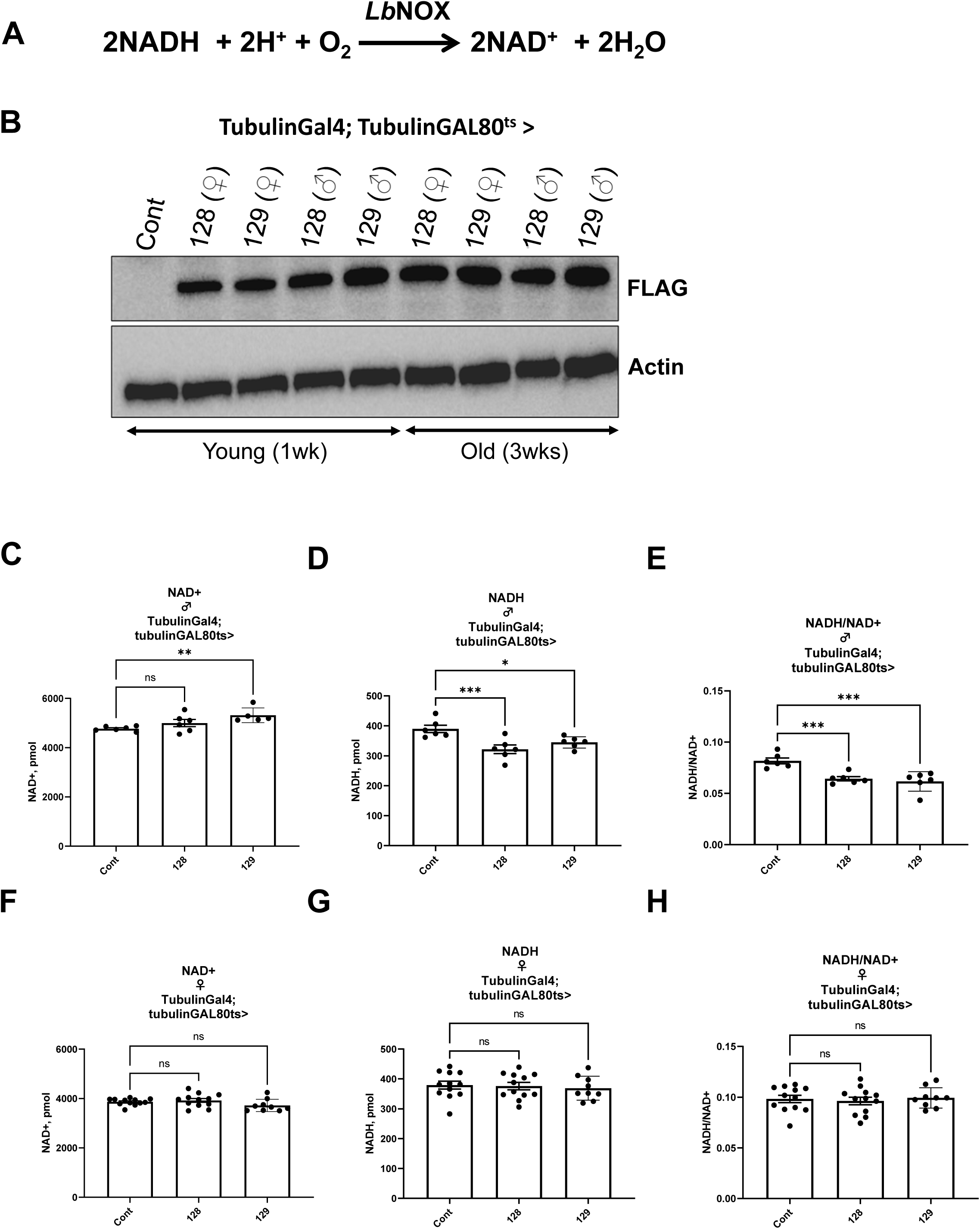
*Lb*NOX expression alters NAD(P)H/NAD(P)^+^ ratios in *Drosophila* in a sex-dependent manner. (**A**) Schematic showing the chemical reaction catalyzed by *Lb*NOX enzyme. (**B**) Representative immunoblot image showing the expression of *Lb*NOX in flies of mentioned ages and genotypes. Levels of NAD^+^ (**C**), NADH (**D**) and NADH/NAD^+^ ratio (**E**) in one week old male flies expressing *Lb*NOX lines 128 and 129. Levels of NAD^+^ (**F**), NADH (**G**) and NADH/NAD^+^ ratio (**H**) in one week old female flies expressing *Lb*NOX lines 128 and 129. Values indicated in (C-H) are from 10 flies. The statistical significance indicated for (C-H) represents a one-way ANOVA followed by uncorrected Fishers Least Significant Difference (LSD) test. The error bars in (C-H) represent mean ± s.d.

We tested the impact of ubiquitous *Lb*NOX expression on NAD^+^ levels and related metabolites *i.e.* NADH, NADP^+^, and NADPH in whole flies. Surprisingly, we observed strikingly different effects on NAD(P)(H) levels and corresponding redox ratios between male and female flies, which reflects sex-specific alterations (**Figure 1C-H**, **Sup. Figure S1A-L**). In male flies of both transgenic lines 128 and 129, we observed a decrease in NADH, which also translated to a robust decrease in NADH/NAD^+^ ratio but with no observed effect on the total NAD pool (NAD^+^ + NADH) (**Figure 1C-E**, **Sup. Figure S1A**). At the same time, in female flies of both transgenic lines 128 and 129, neither NADH levels nor the NADH/NAD^+^ ratio was impacted compared to controls, and no alteration in the total NAD pool was observed (**Figure 1F-H**, **Sup. Figure S1B**).

Interestingly, *Lb*NOX expression in both male and female flies of the transgenic lines 128 and 129 led to a robust increase in NADPH levels (**Sup. Figure S1E, F**) with an increase in NADPH/NADP^+^ ratio only in the female flies of the line 128 (**Sup. Figure S1G, H**). We also observed that the total NADP pool (NADP^+^ + NADPH) was increased in both females and males in transgenic lines 128 and 129 (**Sup. Figure S1I**, **J**). As an illustration, we measured all four coenzymes NAD^+^, NADH, NADP^+^, and NADPH in control male and female flies to highlight that the NAD pool is ∼20 times larger than the NADP pool and that the NADP pool is mostly in the reduced form (**Sup. Figure S1K, L**).

In summary, we have created a novel transgenic xenotopic tool that perturbs NAD(P)(H) metabolites in *Drosophila* in a sex-dependent manner. The observed changes are summarized in **Supplementary Table S1**.

### *Lb*NOX expression rescues neurodegeneration associated with alpha-B crystallin (CryAB)-induced reductive stress in female flies

Reductive stress is defined as an excessive level of reducing equivalents in the form of NADH, NADPH, or GSH (11, 14) or low abundance of reactive oxygen species (ROS) due to inactive electron transport chain (ETC) or hyperactivity of the antioxidant systems (15). Previous studies have described a model of reductive stress in *Drosophila* via expression of the dominant *R120G* mutation in the *aB-crystallin* gene (*CryAB^R120G^*) (16). In humans, the autosomal dominant R120G mutation in the aB-crystallin gene (*CryAB^R120G^*) manifests as adult-onset cataracts, skeletal muscle weakness, and heart failure (17). When mutant CryAB is expressed in *Drosophila* eye, it causes neuronal cell death that can be easily visualized by the appearance of fused glossy disorganized ommatidia under the light microscope (16). The current model of *CryABR^120G^* reductive stress predicts that overabundance of both NADPH and GSH exacerbates the phenotype where lowering cellular NADPH or GSH levels can ameliorate the observed phenotypes seen in flies expressing *CryAB^R120G^* (16). We decided to explore how sex-dependent modulation of NAD(P)(H) levels imposed by *Lb*NOX expression impacts the *CryAB^R120G^* phenotype in *Drosophila. Lb*NOX expression alone did not affect eye morphology in female (**Figure 2A-C**) or male flies (**Figure 2G-I**). As was previously demonstrated, expression of mutant *CryAB^R120G^* resulted in a strong neuronal cell death phenotype in both female (**Figure 2D**) and male flies (**Figure 2J**) characterized by necrotic patches, blebs and fused ommatidia. The phenotype in male flies is relatively drastic with additional reduced eye size and discoloration. Expression of *Lb*NOX transgenic line 128 (but not line 129) completely rescued the neuronal cell death phenotype in female flies (**Figure 2E, F**) while the neuronal cell death phenotype in males was not modified (**Figure 2K**). We note that *Lb*NOX expression boosted NADPH levels in both female lines 128 and 129, while it reliably increased NADPH/NADP^+^ ratio only in line 128, where *Lb*NOX expression completely rescued the *CryAB^R120G^* phenotype. We also note that the *CryAB^R120G^* eye phenotype is much more severe in males compared to females (**Figure 2D, J**). *Lb*NOX expression appeared to rescue the reduced eye size in male flies but did not rescue the neuronal death (**Figure 2K, L**).

**Figure 2:**
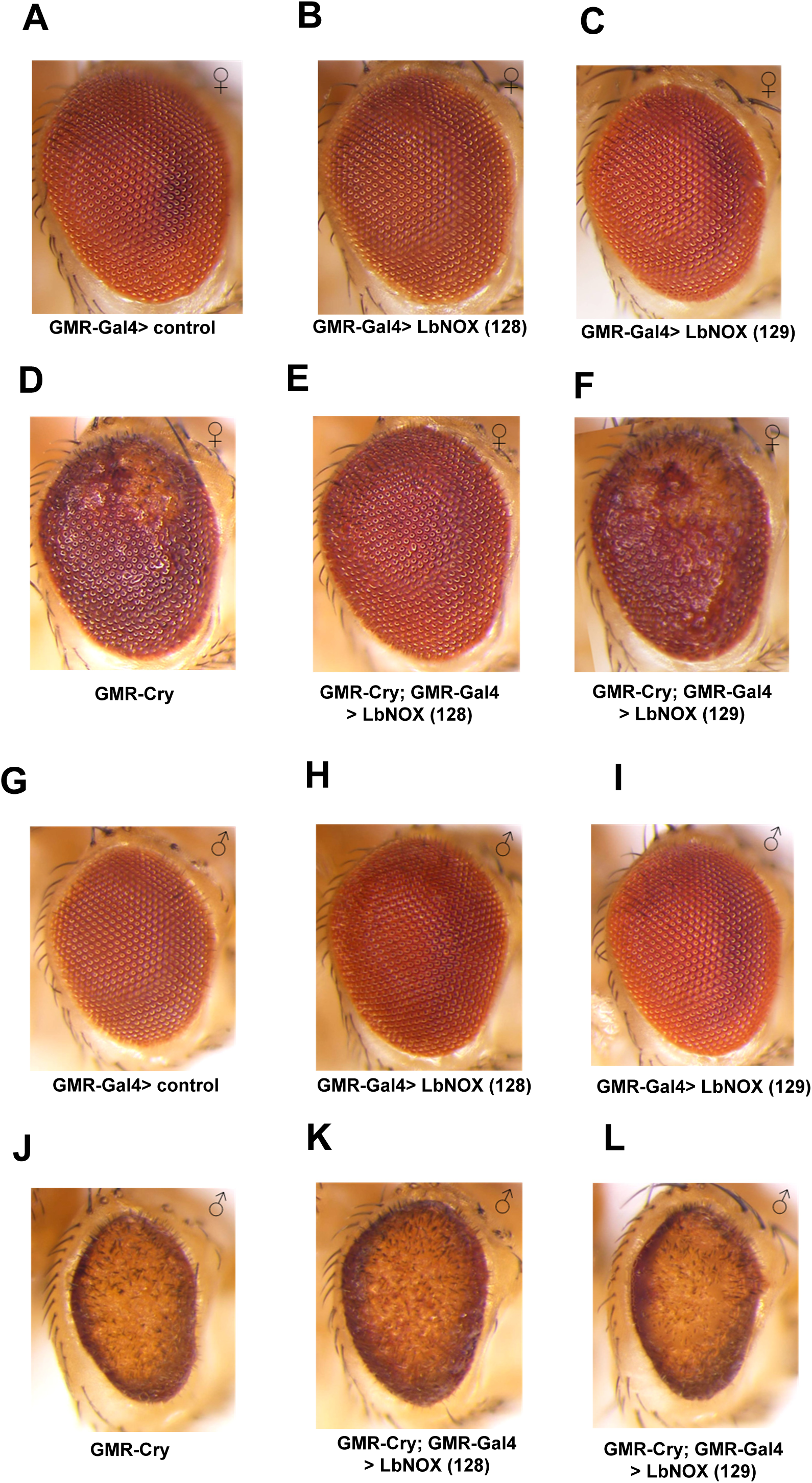
*Lb*NOX expression rescues neurodegeneration associated with alpha-B crystallin (CryAB)-induced reductive stress in female flies in a sex-dependent manner. Representative bright field images of *Drosophila* compound eye in (**A**, **G**) wildtype control (GMR>attp2) female and male flies respectively. (**B**, **H**) flies expressing *Lb*NOX line 128 under GMR-Gal4. (**C**, **I**) flies expressing *Lb*NOX line 129 under GMR-Gal4. (**D**, **J**) flies expressing mutant CryAB under the control of GMR-Gal4. (**E**, **F**, **K**, **L**) flies expressing both mutant CryAB and *Lb*NOX using the mentioned transgenic lines.

Our data suggests that the minor differences observed between the two *Lb*NOX transgenic lines in their effects on total levels of NAD(P)(H) nicotinamide dinucleotides may have a strong impact on the functional outcome as we observed in a model of reductive stress. Notably, female flies had the unique capacity to buffer NADH/NAD^+^ ratio, and despite expression of *Lb*NOX, no decrease in NADH/NAD^+^ ratio was observed exclusively in female *Lb*NOX lines 128 and 129.

### *Lb*NOX expression protects against the detrimental effects of oxidative stress

We further tested how different stress conditions (starvation, oxidative stress, or aging used as a model of integrative stress) affect levels of NAD(P)(H) nicotinamide dinucleotides and their ratios in whole flies and whether *Lb*NOX expression can modify these changes and increase survival of flies under these conditions. We placed 1week-old (1wk old) control female flies on food containing only 1% agar (for inducing starvation stress) or food containing 1% agar with 5% of sucrose and 10 mM of methyl viologen dichloride hydrate (paraquat) (redox cycler that induces oxidative stress) (18). We also aged flies for three weeks at 29°C as an additional stress condition. Both starvation and oxidative stress dramatically reduced NADH levels, as well as the NADH/NAD^+^ ratio, with no effect on the NAD^+^ levels or the total NAD pool (**Figure 3A-C**, **Sup. Figure S2A**). All three stress conditions significantly decreased NADPH levels and the total NADP pool, except for aging, which did not impact the total NADP pool (**Figure 3D-F**, **Sup. Figure S2B**).

**Figure 3:**
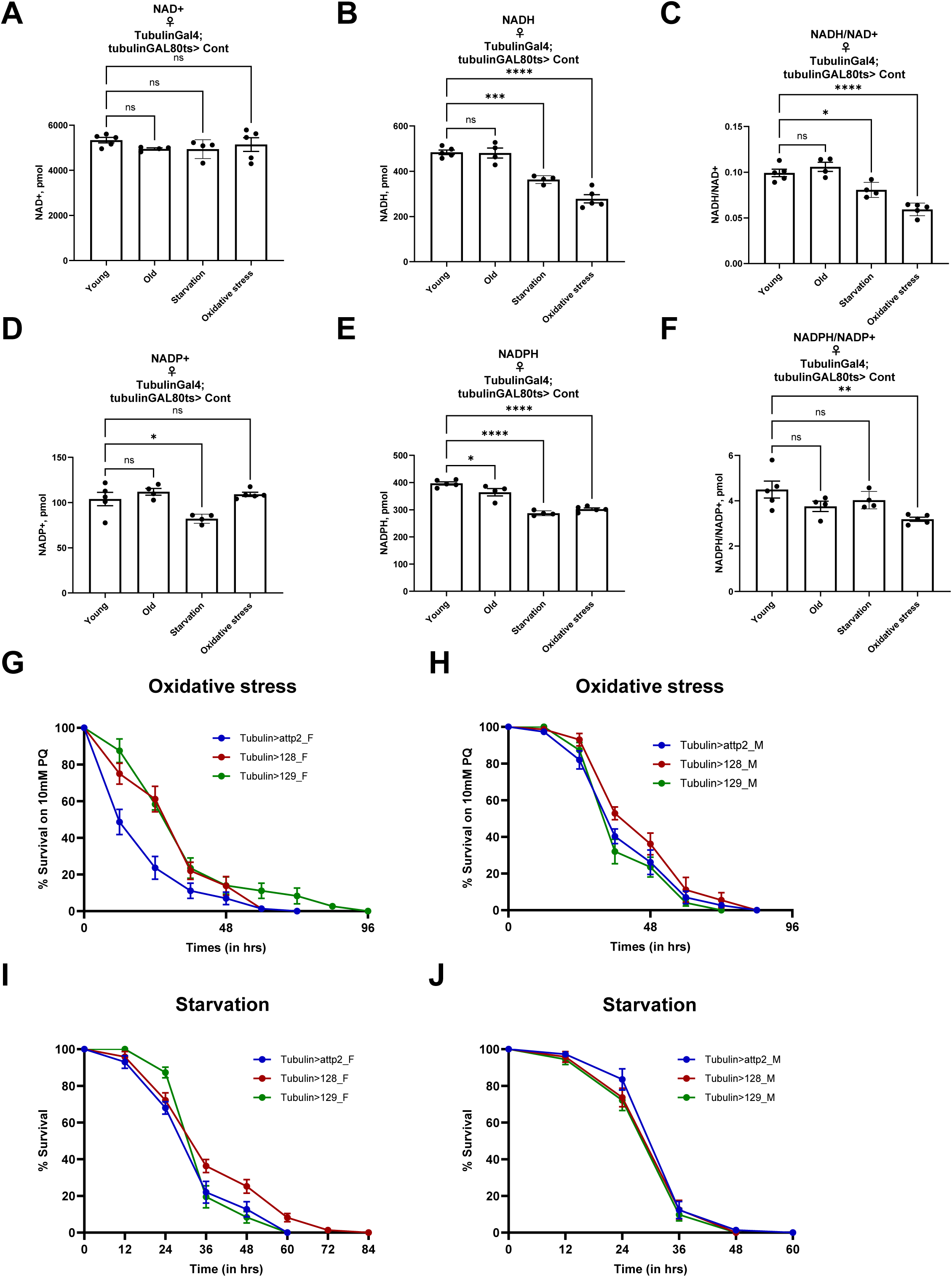
*Lb*NOX expression protects against detrimental effects of oxidative stress. Levels of NAD^+^ (**A**), NADH (**B**), and NADH/NAD^+^ ratios (**C**) in female flies that were 1 week old (young), 3 weeks old (old), subjected to starvation or oxidative stress for 16 hrs. Levels of NADP^+^ (**D**), NADPH (**E**) and NADPH/NADP^+^ ratios (**F**) in female flies that were 1 week old (young), 3 weeks old (old), subjected to starvation or oxidative stress for 16 hrs. Survival of one week old female (**G**) and male (**H**) flies expressing attp2(control) or *Lb*NOX ubiquitously when subjected to oxidative stress. Survival of one week old female (**I**) and male (**J**) flies expressing attp2(control) or *Lb*NOX ubiquitously when subjected to starvation stress. Values indicated in (A-F) are from 10 flies. The statistical significance indicated for (A-F) represents a one-way ANOVA followed by uncorrected Fishers Least Significant Difference (LSD) test. The error bars in (A-F) represent mean ± s.d. The error bars in (G-J) represent mean ± s.e.m.

To test if *Lb*NOX expression restores the decreased NADH and NADPH levels and the decreased NADH/NAD^+^ and NADPH/NADP^+^ ratios in flies under oxidative stress, we placed 1wk old control or *Lb*NOX-expressing female flies on food containing 1% agar with 5% sucrose and 10 mM of paraquat and compared NAD(H) and NADP(H) levels, as well as NADH/NAD^+^ and NADPH/NADP^+^ ratios, to control flies kept on a standard diet. We observed that expression of *Lb*NOX partially rescued the alterations associated with the oxidative stress for both NAD(P)H levels and NAD(P)H/NAD(P)^+^ ratios (**Sup. Figure 2C-F**).

We next explored how *Lb*NOX expression affects the survival of flies under oxidative stress or starvation stress. *Lb*NOX expression significantly improved the survival of female flies under oxidative stress (**Figure 3G**) with a moderate effect in male flies (**Figure 3H**). Under starvation condition, we observed only a minor effect of *Lb*NOX expression on the survival of female flies with no effect on male flies (**Figure 3I, J**).

In summary, we demonstrate that *Lb*NOX expression, despite promoting NADH-consumption, can counteract the detrimental effects of oxidative stress by robustly boosting both NAD(P)H levels and NAD(P)H/NAD(P)^+^ ratios.

### *Lb*NOX expression prolongs *Drosophila* lifespan in a sex-dependent manner

We further sought to evaluate the effects of *Lb*NOX expression on lifespan using both the 128 and 129 lines. To avoid potential contribution from developmental effects and differences in genetic backgrounds, we used the GeneSwitch (GS) inducible Gal4/UAS expression system (19, 20), where UAS-transgene expression is driven by Gal4 when flies are fed mifepristone (RU486). Expression of *Lb*NOX 128 line significantly extended lifespan in female flies using two different ubiquitous GeneSwitch Gal4 drivers: Daughterless-GeneSwitch (Da-GS-Gal4) and Actin-GeneSwitch (Actin-GS-Gal4) (**Figure 4A, C**). *Lb*NOX 128 expression extended lifespan by 17.9% (p < .0001, log-rank test) under Da-GS-Gal4 and by 16.3% (p < .0001, log-rank test) with Actin-GS-Gal4. In males, *Lb*NOX 128 expression did not have a significant effect on lifespan under Da-GS-Gal4 (p = 0.1947, log-rank test) and extended lifespan by 6% under Actin-GS-Gal4 (p = 0.0081, log-rank test) (**Figure 4B, D**). *Lb*NOX 129 expression extended lifespan by 7.7% under Da-GS-Gal4, but the effect was not statistically significant (p = 0.8332, log-rank test) in females, whereas its expression did not affect lifespan in males (**Figure 4E, F**). *Lb*NOX 129 expression extended lifespan by 3.1% under Actin-GS-Gal4, but the effect was not statistically significant (p = 0.5842, log-rank test) in females, while its expression significantly decreased lifespan by 3.1% in males (p = 0.0086, log-rank test) (**Figure 4G, H**).

**Figure 4:**
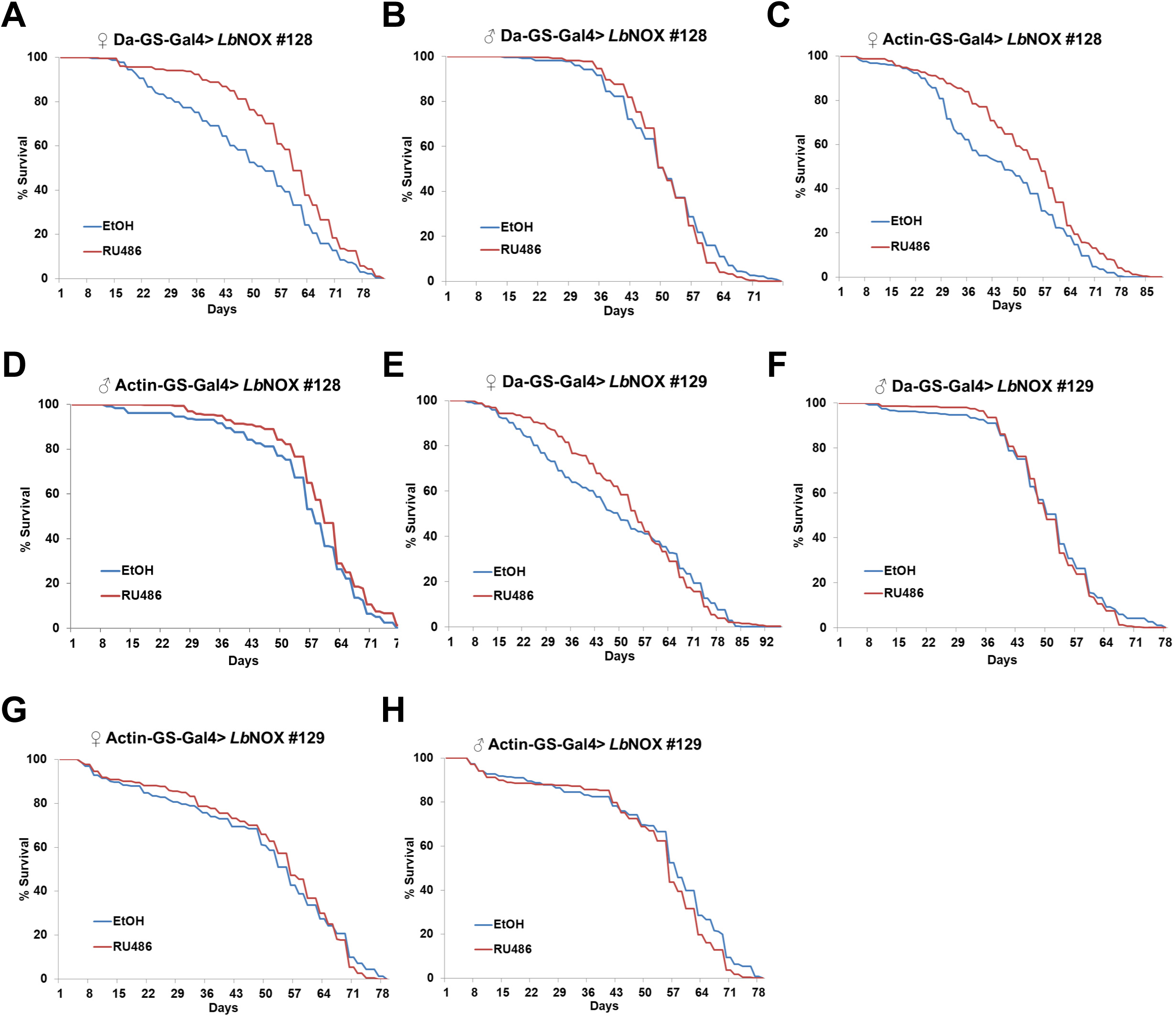
*Lb*NOX expression prolongs *Drosophila* lifespan in a sex-dependent manner. (**A**) Ubiquitous adult-onset expression of *Lb*NOX #128 with Da-GS-Gal4 increases lifespan in females. p < 0.0001. (**B**) Ubiquitous adult-onset expression of *Lb*NOX #128 with Da-GS-Gal4 does not affect lifespan in males. p = 0.1947. (**C**) Ubiquitous adult-onset expression of *Lb*NOX #128 with Actin-GS-Gal4 increases lifespan in females. p < 0.0001. (**D**) Ubiquitous adult-onset expression of *Lb*NOX #128 with Actin-GS-Gal4 increases lifespan in males. p = 0.0081. (**E**) Ubiquitous adult-onset expression of *Lb*NOX #129 with Da-GS-Gal4 does not affect lifespan in females. p = 0.8332. (**F**) Ubiquitous adult-onset expression of *Lb*NOX #129 with Da-GS-Gal4 does not affect lifespan in males. (**G**) Ubiquitous adult-onset expression of *Lb*NOX #129 with Actin-GS-Gal4 does not affect lifespan in females. p = 0.5842. (**H**) Ubiquitous adult-onset expression of *Lb*NOX #129 with Actin-GS-Gal4 decreases lifespan in males. p = 0.0086.

In summary, we demonstrate that ubiquitous *Lb*NOX expression extends *Drosophila* lifespan in a sex-dependent manner.

### NAD precursor supplementations do not further extend lifespan of *Drosophila* expressing *Lb*NOX ubiquitously

We further tested whether the effects of ubiquitous *Lb*NOX expression are comparable to the supplementation of common NAD precursors nicotinamide riboside (NR) and nicotinamide mononucleotide (NMN) based on NAD(P)(H) levels and NAD(P)H/NAD(P)^+^ ratios in whole flies. Supplementation of NMN has previously been shown to significantly rescue the shortened lifespan in a *Drosophila* model of Werner syndrome (21). Similarly, supplementation of NR significantly rescues impaired muscle function in a *Drosophila* model of Barth Syndrome (22). We first supplemented control flies with either NR (100 µM or 1 mM) or NMN (100 µM) for 7 days and measured the levels of NAD(H) and NADP(H) coenzymes in whole flies. In these experiments, we used female flies only, as the beneficial effect of *Lb*NOX expression in the model of reductive stress was observed exclusively in females. Supplementation with either NR or NMN increased levels of NAD^+^ as well as the total NAD pool and promoted a robust pro-oxidative shift (a decrease) in NADH/NAD^+^ ratio (**Figure 5A-C**, **Sup. Figure S3A**). In contrast, only supplementation with NMN increased NADP^+^ levels (with no significant effect on the total NADP pool) and decreased NADPH/NADP^+^ ratio (**Figure 5D-F**, **Sup. Figure S3B**). We did not observe any significant difference between the two NR concentrations tested.

**Figure 5:**
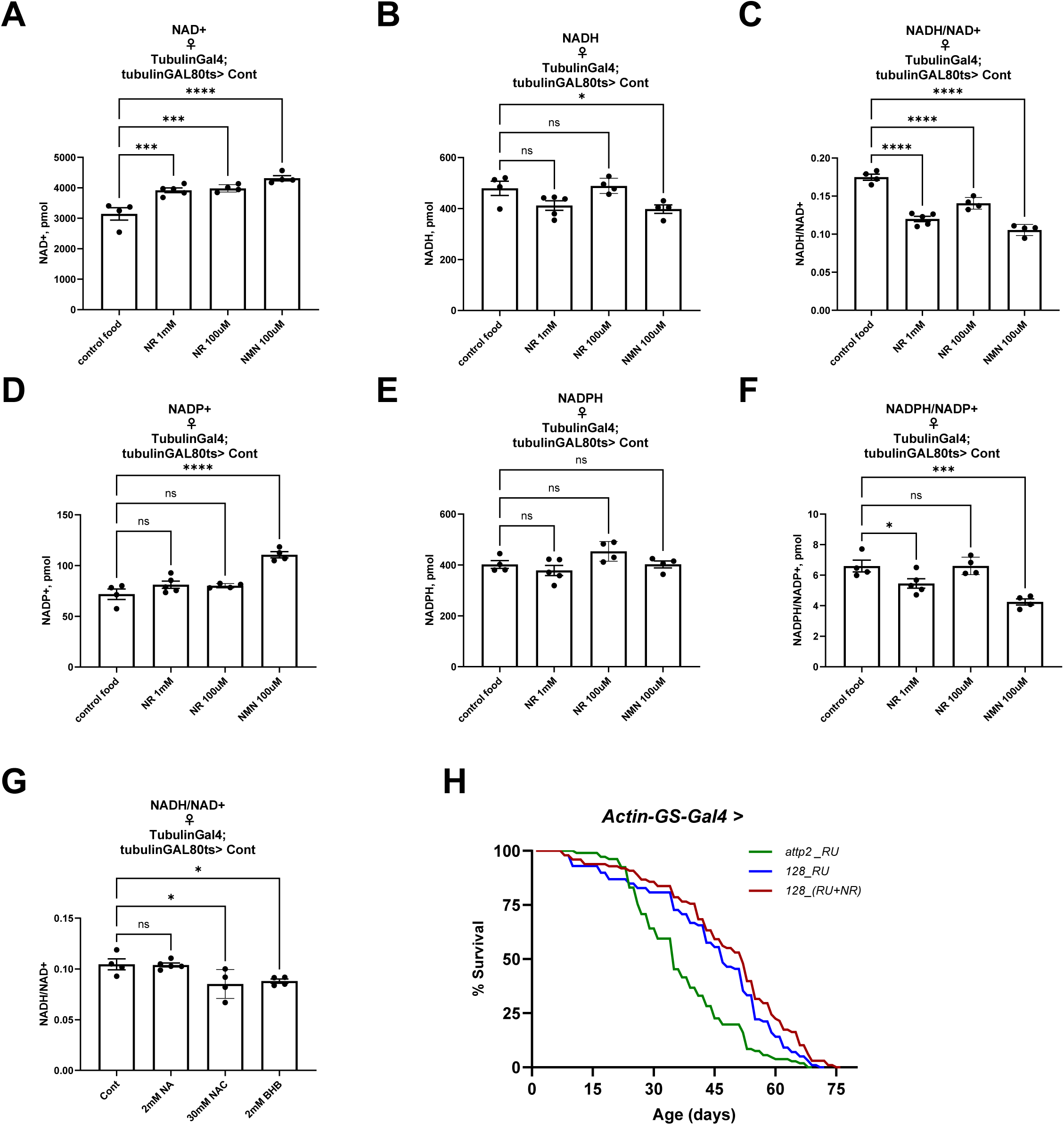
NAD precursor supplementations do not further extend lifespan of *Drosophila* expressing *Lb*NOX ubiquitously. Levels of NAD^+^ (**A**), NADH (**B**) and NADH/NAD^+^ ratios (**C**) in female control flies fed with food containing 1mM NR, 100μM NR or 100 μM NMN compared to flies fed on control food. Levels of NADP^+^ (**D**), NADPH (**E**) and NADPH/NADP^+^ ratios (**F**) in female control flies fed with food with 1mM NR, 100μM NR or 100 μM NMN. (**G**) The NADH/NAD^+^ ratio in female control flies fed with food containing 2mM NA, 30mM NAC or 2mM BHB compared to flies fed on control food. (**H**) Ubiquitous adult-onset expression of *Lb*NOX #128 with Actin-GS-Gal4 increases lifespan in females that is not further extended with 0.5 mM NR supplementation. Values indicated in (A-G) are from 10 flies. The statistical significance indicated for (A-G) represents a one-way ANOVA followed by uncorrected Fishers Least Significant Difference (LSD) test. The error bars in (A-G) represent mean ± s.d.

We further tested how supplementation of nicotinic acid (NA), ²-hydroxybutyrate (BHB), and N-acetyl-cysteine (NAC) affected levels of NAD(H) and NADP(H) coenzymes in whole flies. NA has previously been shown to increase the intracellular NAD levels in Hs68 cells and *C. elegans,* as well as to extend worm’s lifespan (23). In *Drosophila*, expression of *Drosophila* nicotinamidase (D-NAAM/NAMase) converts nicotinamide to NA, which decreases NADH/NAD^+^ levels and extends lifespan (24). BHB is a circulating ketone body, which is metabolized into acetoacetate by ²-hydroxybutyrate dehydrogenase 1 (BDH1) in mitochondria of target tissues, thereby generating NADH. In humans, food supplementation with ²-hydroxybutyrate or sodium butyrate has been associated with multiple health benefits (25). Finally, NAC is associated with lifespan benefits across different species (1) and may indirectly affect NAD(P) pools by acting as an antioxidant. We found that supplementation of 2 mM NA did not affect NADH/NAD^+^ or NADPH/NADP^+^ ratios in *Drosophila* (**Figure 5G**, **Sup. Figure S3C-G**). However, supplementation of 30 mM NAC or 2 mM BHB significantly (but less potently than NR or NMN) decreased NADH/NAD^+^ without impacting the NADPH/NADP^+^ ratio (**Figure 5G**, **Sup. Figure S3C-G**).

As NR significantly decreased NADH/NAD^+^ ratio (**Figure 5C**) and *Lb*NOX expression significantly increased the NADP pool (**Sup. Figure S1J**) and prolonged lifespan (**Figure 4C**) in female flies, we further tested if the combination of NR supplementation and *Lb*NOX expression would be more efficient than *Lb*NOX expression alone. In female flies, supplementation of 0.5 mM of NR along with *Lb*NOX expression did not significantly extend lifespan compared to *Lb*NOX expression alone (**Figure 5H**).

In summary, we assessed how different NAD boosters or modulators (NR, NMN, NA, BHB, and NAC) affect NAD metabolism in *Drosophila* and demonstrated that the lifespan extension achieved via *Lb*NOX expression cannot be further extended with NR supplementation.

### *Lb*NOX expression reprograms *Drosophila* metabolism with a strong impact on *de novo* NAD biosynthesis

To better understand the mechanism of lifespan extension by *Lb*NOX expression, we performed metabolomic profiling on either control flies or two different *Lb*NOX expressing lines under the control of the ubiquitous temperature sensitive (*tubulin-Gal4, tubulin-Gal80^ts^*) driver after 7 days of expression (**Figure 6A-C**, **Supplementary Table S2**). Principal component analysis (PCA) of the measured metabolites clearly distinguished control flies from *Lb*NOX lines 128 and 129 (**Sup. Figure S4A**). We identified 70 metabolites were significantly altered in the *Lb*NOX 128 line and 19 metabolites were significantly altered in the *Lb*NOX 129 line; while 18 metabolites were significantly and commonly changed in both *Lb*NOX lines (**Figure 6D**). To understand which metabolic pathways were among the most significantly changed by *Lb*NOX expression, we applied Metabolite Set Enrichment Analysis (MSEA). We found that the TCA cycle and tryptophan metabolism were among the metabolic pathways most affected by *Lb*NOX expression (**Figure 6E-F**, **Supplementary Table S3**). Interestingly, *Lb*NOX expression dramatically altered the levels of several metabolites (tryptophan, kynurenine, kynurenic acid, quinaldic acid, tryptamine, indole-3-acetic acid, indole-3-carboxylic acid, and indoleacetic acid) in the tryptophan metabolic pathway that are involved in *de novo* NAD biosynthesis (**Figure 6G**, **H**). We also noted that *Lb*NOX expression in both lines 128 and 129 robustly decreased levels of metabolites attributed to the indole pyruvate pathway, which is typically linked to tryptophan and kynurenine metabolism (26–30). The results of our metabolomic profiling may explain the stronger effects of *Lb*NOX expression on cellular metabolism in the 128 line. In addition, increase in the NADP pool under *Lb*NOX expression may explain the upregulation of NAD *de novo* biosynthesis and depletion of metabolites in the tryptophan metabolic pathway.

**Figure 6:**
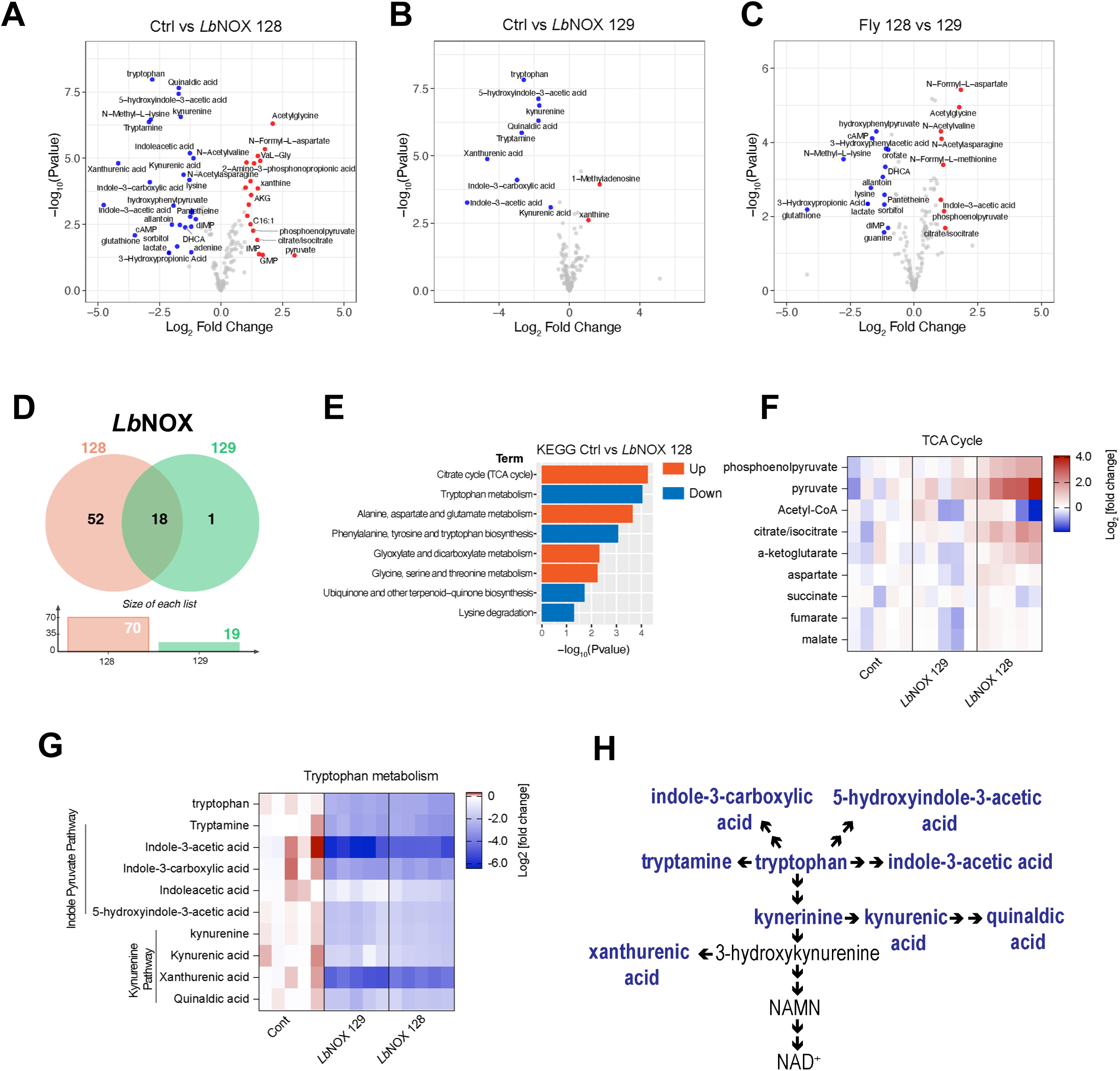
*LbNOX* expression reprograms *Drosophila* metabolism with a strong impact on *de novo* NAD biosynthesis. Volcano plots of targeted metabolomics of female *Drosophila Lb*NOX lines 128 (**A**) and 129 (**B**) relative to controls as well as comparison between 128 and 129 lines (**C**). Accumulated metabolites are shown in red dots, decreased metabolites are shown in blue dots and gray dots represent statistically non-significant changes. The statistical significance for (A-C) represents p value cutoff = 0.05, fold change cutoff = 0.5. (**D**) Venn diagrams of metabolites significantly altered in *Lb*NOX expression lines 128 and 129. (**E**) Kyoto Encyclopedia of Genes and Genomes (KEGG) pathways analysis based on MetaboAnalyst platform for metabolites with *Lb*NOX expression line 128 relative to controls. Orange and blue depict pathways that are upregulated or downregulated, respectively (based on the abundance of the corresponding metabolites). Heatmaps of the most impacted metabolites of TCA cycle (**F**) and tryptophan metabolism and indole pyruvate pathway (**G**) in female *Lb*NOX lines 128 and 129. In heatmaps (F-G) each column represents a biologically independent sample. (**H**) A simplified schematic of the kynurenine and indole pyruvate pathways in *Drosophila*. Two consecutive arrows assume multiple reaction steps. Metabolites from the heatmap (G) are in blue color.

### Tissue-specific *Lb*NOX expression efficiently protects against oxidative stress and rejuvenates sleep profiles in aged flies back to a youthful state

Ubiquitous expression of *Lb*NOX improved the survival of female flies under oxidative stress and had a minor effect on the survival of flies of both sexes under starvation. We further used our genetically encoded tool to test if expression of *Lb*NOX in one tissue alone is sufficient to have an effect similar to that of ubiquitous expression. We expressed *Lb*NOX in adults in neuronal tissues using the *Elav-Gal4, tubulin-Gal80^ts^* temperature-inducible system or in the muscles using the *Dmef-Gal4, tubulin-Gal80^ts^* temperature-inducible system. Surprisingly, we observed that muscle-specific *Lb*NOX expression was much more effective in protecting against oxidative stress in females with no impact on male flies (**Figure 7A, B**), while expression of *Lb*NOX in neurons slightly improved stress tolerance in females but was detrimental in males (**Figure 7C, D**). Both muscle-and neuron-specific *Lb*NOX expression (#128 line) promoted survival of female flies under starvation (**Sup. Figure S5A, C)**.

**Figure 7:**
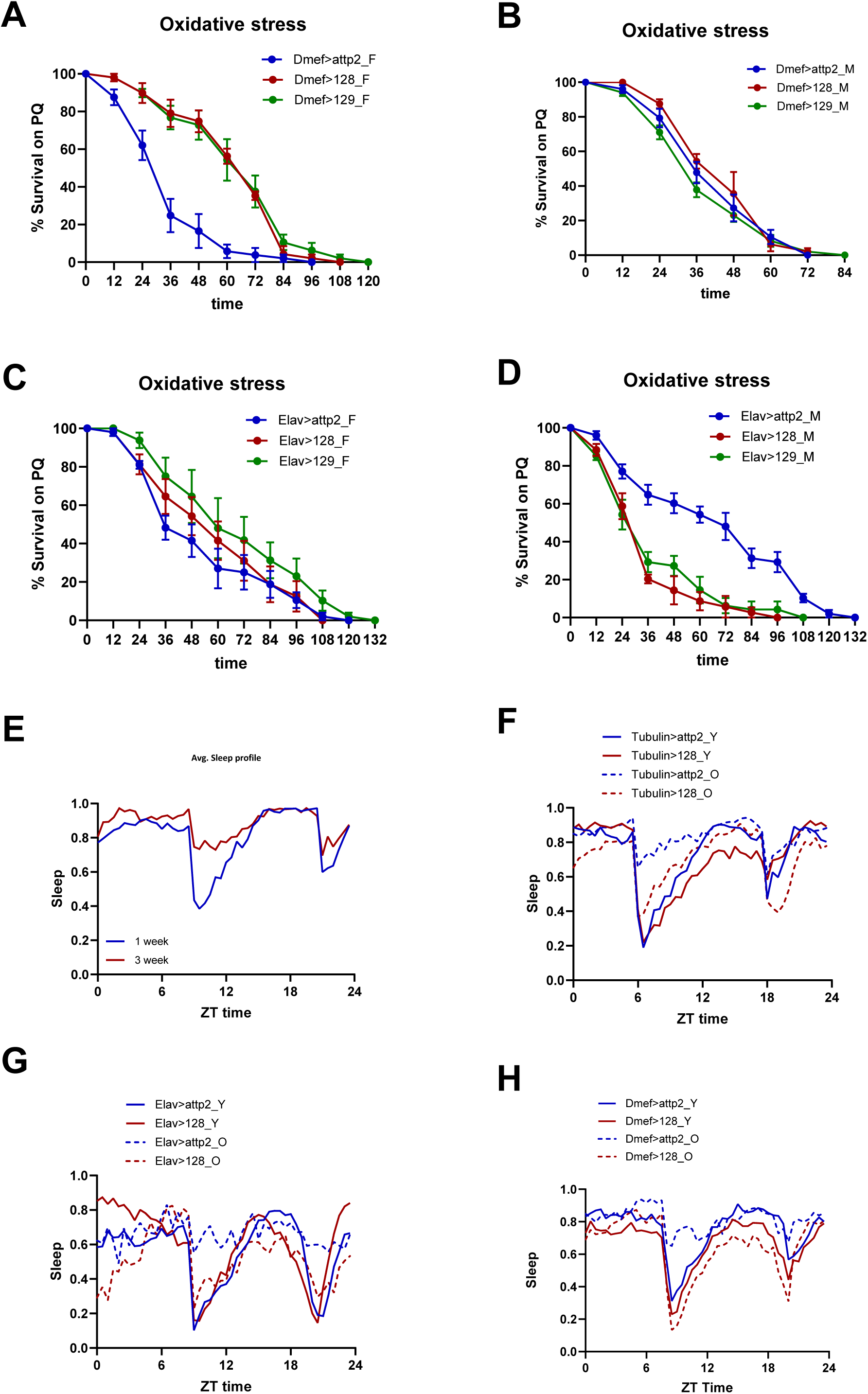
Tissue-specific *Lb*NOX expression efficiently protects against oxidative stress and rejuvenates sleep profiles in aged flies back to a youthful state. Survival of female (**A**) and male (**B**) control flies and flies with muscle-specific *Lb*NOX expression under oxidative stress. Survival of female (**C**) and male (**D**) control flies and flies with neuron-specific *Lb*NOX expression under oxidative stress. Average sleep profile (**E**) of control female flies at 1-week (marked as Y) and 3-weeks (marked as O) of age. ZT time refers to the Zeitgeber time with respect to the onset of light. Average sleep profile **(F)** of control flies or flies with ubiquitous expression of LbNOX #128 at 1-week (marked as Y) and 3-weeks (marked as O) of age. Average sleep profile **(G)** of control flies or flies with neuronal expression of LbNOX #128 at 1-week (marked as Y) and 3-weeks (marked as O) of age. Average sleep profile **(H)** of control flies or flies with muscle-specific expression of LbNOX #128 at 1-week (marked as Y) and 3-weeks (marked as O) of age.

Recent studies have shown the importance of NAD(P) metabolism in the regulation of sleep in *Drosophila* (31, 32), mice (33, 34), and humans (35, 36). As the key genes regulating circadian rhythm have been identified (37) and are conserved from flies to humans (38, 39), *Lb*NOX represents a unique tool to dissect the role of NAD(P) metabolism across different tissues and populations of cells. Sleep is defined in *Drosophila* as a prolonged period of immobility longer than five minutes (40) and can be measured using the *Drosophila* Activity Monitors (DAM) by determining the number and duration of sleep bouts. We first monitored sleep in control female flies aged from 1-week through 5-weeks. We found that aging is associated with disruption of sleep parameters namely sleep bout length and number. The changes start to appear 3-weeks onwards and are maintained thereafter (**Sup. Figure 5E, F**). We found that the sleep bout length increased with age while the sleep bout number decreased with age (**Sup. Figure 5E, F**). The average sleep profile of 3-weeks old fly was significantly altered compared to that of 1-week old fly (**Figure 7E**). We tested if whole-body *Lb*NOX expression could rescue the aging associated disruption of sleep profile. We found that ubiquitous expression of *Lb*NOX line 128 significantly rescued the average sleep profile in 3-weeks old flies (**Figure 7F**) and it also rescued the disrupted sleep bout length and number in old flies (**Sup. Figure 5G, H)**. Sleep is known to be regulated by neuronal inputs hence we tested if a neuron-specific *Lb*NOX expression could rescue the aging associated sleep deficits. We expressed with *Lb*NOX using the Elav-Gal4 driver and intriguingly it rescued the disrupted sleep profile in 3-weeks old flies (**Figure 7G**). The beneficial effects observed with muscle-specific expression of *Lb*NOX on fly’s lifespan and ability to tolerate stress prompted us to further test if muscle-specific expression of *Lb*NOX could exert beneficial effects on sleep as well. Interestingly, the expression of *Lb*NOX using *Dmef-Gal4* driver completely rescued the sleep profile in old flies (**Figure 7H**) demonstrating that tissues-specific modulation of NAD metabolism is sufficient to drive beneficial effects at the organismal level.

In summary, we demonstrate that utilization of *Lb*NOX allows assessing the tissue-specific roles of NAD metabolism in different cellular processes and its impact on overall health-and lifespan of *Drosophila*.

## Discussion

### Exploiting xenotopic tools to manipulate metabolism

In recent years, manipulations of over a hundred different metabolic enzymes have been shown to prolong lifespan in different organisms (1). However, targeting only species-specific enzymes may not be sufficient to reverse age-dependent changes in a particular metabolic pathway and will only partially reverse age-dependent processes. An alternative approach to targeting an organism’s own enzymes is to bring a novel function from other species (a “xenotopic approach”) that would resolve or reprogram a metabolic pathway in a beneficial way for an organism to achieve healthy aging. The yeast *S. cerevisiae* NDI1 and the sea-squirt *Ciona intestinalis* AOX are probably among the most commonly used xenotopic tools in *Drosophila* to study the role of the mitochondrial electron transport chain (ETC) in aging and aging-related diseases (41–44). Alternative yeast NADH dehydrogenase NDI (NADH dehydrogenase internal, faces matrix) can bypass mitochondrial complex I (CI), and alternative ubiquinol oxidase (AOX) can bypass complexes III and IV (CIII, CIV). We have previously created and tested several additional xenotopic tools including Methioninase (45), *Lb*NOX (10), TPNOX (46), and *Ec*STH (47). We created transgenic flies expressing Methioninase, a bacterial enzyme capable of degrading methionine to ammonia, a-ketobutyrate, and methanthiol, and demonstrated that either whole-body or tissue-specific Methioninase expression can dramatically extend *Drosophila* health-and lifespan and exert physiological effects associated with methionine restriction (45). Here, we created transgenic flies carrying the *Lb*NOX tool and demonstrated its utility in studying NAD(P) metabolism *in vivo*.

### *Lactobacillus brevis* water-forming NADH oxidase (*Lb*NOX)

Unlike other bacterial NADH oxidases, *Lactobacillus brevis* water-forming NADH oxidase (*Lb*NOX) does not generate H_2_O_2_ and is strictly NADH-specific (**Figure 1A**). In our original study, *Lb*NOX was expressed in HeLa cells (both untargeted (*Lb*NOX) and mitochondria-targeted (mito*Lb*NOX) versions) to demonstrate that direct NAD^+^ recycling is sufficient to restore proliferation in cells with impaired ETC activity (10). Currently, *Lb*NOX and mito*Lb*NOX are commonly used tools to dissipate cytoplasmic or mitochondrial NADH pools (with a concomitant decrease in NADH/NAD^+^ ratio) in settings of both dysfunctional and intact mitochondrial ETC. Due to technical limitations, we were unable to generate transgenic flies expressing the mito*Lb*NOX version. Interestingly, in males, *Lb*NOX expression resulted in a decrease in NADH levels and a concomitant decrease in NADH/NAD^+^ ratio; while in both males and females, expression of *Lb*NOX resulted in both a robust increase in NADPH and the NADPH/NADP^+^ ratio (**Figure 1D, E**, **Sup. Figure S1H**, **I**). These findings highlight the importance of testing xenotopic tools *in vivo* as results previously obtained in mammalian cell culture cannot be extrapolated to model organisms.

Moreover, *Lb*NOX can be coupled with other xenotopic tools targeting redox stress (TPNOX (46) or *Ec*STH (47, 48)) to obtain different outcomes in NADH/NAD^+^ and NADPH/NADP^+^ ratios. Previously, we used *Drosophila* to create a complex genetic background that allowed for the simultaneous targeting of several cancer-related genes (49). As we have limited knowledge of tissue-specific metabolic dysregulation with aging (50) and how different interventions interact in extension of health-and lifespan (51), a similar approach can be applied to combine different xenotopic tools targeting redox stress (or other processes) to study these combinations and their effects on lifespan *in vivo*.

### Targeting NAD metabolism to delay aging

Multiple studies in different organisms have demonstrated a decline in the total levels of NAD^+^ with age in a tissue-specific manner. This has led to the hypothesis that a decline in NAD^+^ levels with age is one of the major contributing factors to the aging process (2–4). Multiple pre-clinical studies in rodents have provided evidence for the beneficial effects of supplementation with NAD biosynthetic precursors that boost total cellular NAD levels (7–9). However, supplementation studies neither allow testing of the tissue-specific roles of restoring NAD^+^ levels nor do they allow investigation of how NADH/NAD^+^ ratio itself regulates the aging process. Moreover, in a systematic review of aging-related NAD changes across different organisms, the authors noted that there are no published papers that address whether NAD^+^ levels decline with age in *Drosophila* model (7). Despite this, targeting NAD metabolism in *Drosophila* via whole-body or neuron-specific expression of *Drosophila* nicotinamidase (D-NAAM/NAMase) increased NAD^+^/NADH levels and extended lifespan (24).

By employing *Lb*NOX as a genetically encoded xenotopic tool (10) and a binary UAS/GAL4 system (52), we were able to test whether directly manipulating NADH/NAD^+^ ratio ubiquitously (mimicking supplementation of NAD precursors) or in a tissue-specific manner affects *Drosophila* aging. We tested how NAD levels change with age and in response to stress, identifying whether *Lb*NOX expression phenocopies supplementation of NAD precursors and whether altering NADH/NAD^+^ ratio alone or in combination with NAD precursors prolongs lifespan. Unexpectedly, we found that NADH/NAD^+^ and NADPH/NADP^+^ ratios were significantly altered under stress conditions, but not with aging (**Figure 3C, F**). This finding agrees with a recent study demonstrating that NADH/NAD^+^ and NADPH/NADP^+^ ratios do not change in mice with healthy aging because they are buffered by the activities of fatty acid desaturases that promote NAD^+^ recycling activity (12). Accordingly, with an oxidative stress-induced decrease in NADH/NAD^+^ and NADPH/NADP^+^ ratios, we demonstrate a protective role of *Lb*NOX expression which robustly increases both ratios despite the NADH-consuming activity of the enzyme under stress conditions (**Sup. Figure S2D, F**). Interestingly, although we do not observe age-induced changes in NADH/NAD^+^ and NADPH/NADP^+^ ratios, we did observe the beneficial effects of *Lb*NOX expression on lifespan (**Figure 4A, C**). There could be several explanations for these beneficial effects. First, age-induced changes in NADH/NAD^+^ and NADPH/NADP^+^ ratios are not homogeneous across different tissues and may not be unidirectional. McReynolds *et al.* employed isotope tracing and mass spectrometry to probe age-related changes in NAD metabolism across tissues in aged mice (53). They found that the decline in NAD with healthy aging is relatively subtle and tissue-dependent (for example the reduced forms of NADH and NADPH significantly declined with age in the liver and increased in the brain) (53). This will result in the absence of aging-induced whole-body changes, while NAD(P) levels can be altered dramatically at the tissue-specific level. Second, age-induced changes in NAD(P) metabolism can be compensated by other metabolic pathways (12). Lien *et al.* observed that levels of NAD^+^, NADH, and the NADH/NAD^+^ ratio were unchanged in plasma, liver, gastrocnemius muscle, and brain tissues with healthy aging; however, aging tissues, particularly the brain, exhibited evidence of up-regulated fatty acid and sphingolipid metabolism reactions that regenerate NAD^+^ from NADH (12).

In addition to testing tissue-specific effects of NAD(P) metabolism, the application of *Lb*NOX as a tool combines two different strategies for maximizing the benefits of extending lifespan: *(i)* supplementation with NAD precursors (such NR and NMN) and *(ii)* directly altering NADH/NAD^+^ and NADPH/NADP^+^ ratios. By testing this strategy in *Drosophila*, we found that combining ubiquitous *Lb*NOX expression with the administration of NR in food did not result in an extended lifespan. We have tested a variety of other NAD boosting compounds and found that NR and NMN were among the most effective. In line with this, Rimal *et al.* discovered that genetic or pharmacological inhibition of reverse electron transfer at mitochondrial Complex I decreased NADH/NAD^+^ ratio and extended *Drosophila* lifespan (54). Testing different strategies will help identify the most effective combination for potential human translation to forestall aging.

### Rebalancing of reducing equivalents between NAD and NADP pools

The NAD and NADP pools are interconnected via mitochondrial transhydrogenase (NNT) and multiple dehydrogenases in different cellular compartments with dual NAD(P) specificity (46, 55, 56). In our study, we also found that expression of the NADH-consuming enzyme (*i.e. Lb*NOX) impacts both NAD(H) and NADP(H) metabolites. We observed that *Lb*NOX expression can increase NADP(H) levels and NADPH/NADP^+^ ratio. Our findings are in agreement with recent studies that identify NADPH regeneration as an important metabolic parameter under ETC inhibition and elevated NADH/NAD^+^ (57–59). Therefore, in principle, a pro-oxidative shift (a decrease) in NADH/NAD^+^ ratio promoted by *Lb*NOX can boost NADPH levels. Notably, in our measurements of NAD(P)(H) metabolites, we use enzymatic cycling and whole-body fly homogenates. Further studies are needed to better understand the role of compartment-specific changes in both NADH/NAD^+^ and NADPH/NADP^+^ using genetically encoded sensors in healthy aging as well as under various stress conditions. In flies (as well as in humans) the only route to produce NADP from NAD is NAD kinase (both cytosolic and mitochondrial versions exist) but surprisingly very little attention has been given to NADK regulation in the context of aging (60). One promising future direction will be to use knockout flies in concert with genetically encoded tools *Lb*NOX, TPNOX, or *Ec*STH to dissociate NAD metabolism from NADP metabolism as well as to uncouple the regulation of NADP(H) production from cellular metabolism linked to ROS.

### Sleep

Sleep disturbances are observed with aging and may serve as a risk factor for developing cognitive impairment and Alzheimer’s disease (61–63). *Drosophila* is a powerful genetic model system for studying sleep (64). Previous studies have established the importance of NAD(P) metabolism in the regulation of sleep in *Drosophila* (31, 32), mice (33, 34), and humans (35, 36). Using LbNOX allows us to test the involvement of NAD(P) metabolism in different tissues in the regulation of sleep. Excitingly, we find that perturbing NAD(P) metabolism in non-neuronal cells is sufficient to rejuvenate sleep profiles to a youthful stage. Lane *et al.* found genetic loci for self-reported insomnia symptoms encompassing genes expressed in skeletal muscle (65). Using LbNOX or other xenotopic tools will allow us to dissect non-neuronal mechanisms of sleep regulation.

### Sex-specific effects

The regulation of aging-related processes is often sex-specific and targeting of a multitude of metabolic pathways often delays aging in one sex but not in another (1). In agreement with this, we observed marked sex-dependent effects of *Lb*NOX expression in *Drosophila*. These sex-specific effects of alterations in NAD metabolites are also observed in healthy humans, as women have higher plasma NAD^+^/NADH ratios than men (66). Accordingly, Van der Velpen *et al.* demonstrated sex-specific alterations in the levels of intermediates in NAD metabolism comparing wildtype (as well as AD-relevant) mice of different sexes (67). We observed that male and female flies have drastically different phenotypes under *Lb*NOX expression. In general, *Lb*NOX expression has stronger protective effects against different stresses and in extending lifespan in females than in males. In *Drosophila*, sex determination is achieved in a cell-specific manner by a balance of female determinants on the X chromosome and male determinants on the autosomes. As we observe that *Lb*NOX expression just in one tissue has a significant effect on the whole organismal physiology in a sex-specific manner; in the future, we can use available *Drosophila* tools together with *Lb*NOX to alter sex only in specific tissues to determine if it would reverse these *Lb*NOX-associated phenotypes. This would allow studying interactions between NAD metabolism and sex-specific effects on a tissue-specific level.

Overexpression of mutated human alpha-B crystallin (CryAB^R120G^) was previously used to model reductive stress in both mice and flies (16, 68). It is believed that CryAB^R120G^ overexpression leads to protein aggregation which triggers NRF2 activation and upregulation of antioxidant machinery, ultimately leading to an elevated GSH/GSSG ratio (11). Xie *et al.* demonstrated that CryAB^R120G^ expression in *Drosophila* mounts impaired cardiac function when expressed in the heart or severely impaired eye development when expressed in the eye (16). The authors concluded that CryAB^R120G^ expression in *Drosophila* leads to reductive stress, at least in part through an elevated GSH/GSSG ratio by demonstrating that RNAi-mediated knockdowns of multiple NADPH-producing enzymes (G6PD, PGD, IDH, MEN) suppress this CryAB^R120G^ pathology (16). Paradoxically, our study shows that the eye phenotype is only rescued by *Lb*NOX expression line 128, in which *Lb*NOX expression leads to an increase in NADPH levels and NADPH/NADP^+^ ratio (**Figure 2E**, **Supp. Figure S1F, H**). We also note that the major difference between males and females expressing *Lb*NOX is that in males we observe a clear decrease in the total NADH/NAD^+^ ratio, while in females this is not the case (**Figure 1E, H**). Reductive stress is a relatively new concept and the exact mechanism by which reductive stress contributes to impaired cardiac function or eye pathology is unknown (16). Nonetheless, our results suggest a useful paradigm to study this phenomenon in more depth.

## Materials and Methods

### *Drosophila* stocks and maintenance

Flies (*Drosophila melanogaster*) were fed a standard cornmeal medium containing yeast, dextrose, sucrose, and molasses and reared at 25°C in incubators with a 12-hour light-dark cycle. For all experiments, the closest genetic control flies (used to generate the transgenic line) were used and treated the same way as the experimental flies. The list of fly stocks used in the study is detailed in the Key Resource Table (**Supplementary Table S4**).

### Acquisition of eye pictures

All the flies were collected within 24hrs of eclosion and were aged for 5 days at 25°C followed by snap-freezing on dry ice. For each genotype, five-six individual fly eyes were imaged using Nikon SMZ1270 microscope at different focal lengths and were combined later using the freely available CombineZP software.

### Lifespan Analysis

Flies were collected within 24-48 hrs from eclosion and allowed to mate for 1-2 days followed by sorting by sex under CO_2_ anesthesia. Post separations flies were reared at standard density (20-25 flies per vial) at 25°C (unless otherwise mentioned) and 60% humidity with 12 hrs On/Off light cycle. Flies were flipped onto fresh vials every two days, and the dead flies were counted. For flies carrying GeneSwitch element (GS lines), gene expression was induced by feeding flies on fly food containing the chemical RU-486 (Cayman chemicals, Item No. 10006317) dissolved in ethanol at a final concentration of 200 uM (86ug/ml). The control flies were reared on food containing equal amounts of the vehicle/solvent (ethanol).

### Starvation assay

All the flies were allowed to mate for 1-2 days post eclosion followed by separation by sex and were reared at appropriate temperature for one week or 3 weeks as referred in the text. For induction of starvation stress the separated flies were placed on vials having freshly prepared 1 % agar. The total fly density per vial was kept between 12-15 flies and the number of dead flies was counted twice a day (morning and evening) until all flies died.

### Oxidative stress assay

All the flies were allowed to mate for 1-2 days post eclosion followed by separation by sex and were reared at appropriate temperature for one week or 3 weeks as referred in the text. For induction of oxidative stress, the flies were placed on freshly prepared food containing 1% agar, 5% sucrose, and 10mM of paraquat. The total fly density per vial was kept between 12-15 flies and the number of dead flies was counted twice a day (morning and evening) until all died.

### Quantification of NAD^+^, NADH, NADP^+^ and NADPH

Ten flies were placed into Precellys Hard Tissue Homogenizing Kit 2 mL Reinforced Tubes (Precellys #: P000916-LYSK0-A.0) and immediately 600 μL of wet ice-cold 1:1 mixture of PBS and 1% dodecyl trimethylammonium bromide (DTAB) in 0.2 M NaOH was added. Flies were homogenized using Bertin Precellys 24 Lysis Biological Tissue Homogenizer. Immediately after homogenization step tubes were centrifuged at 21000 rcf speed for 10 min to pellet down debris and two 200 μL aliquots were used to process oxidized and reduced nucleotides as previously described(46) with minor modifications indicated below. When transferred to all-white 96-well plates samples for NADH and NAD^+^ determination were diluted 5 times with the dilution buffer (equal volumes of PBS, base solution with 1% DTAB, 0.4N HCl, 0.5M Trizma) containing 10 μM ascorbic acid (10 μL sample + 40 μL of the dilution buffer). This dilution is critical to maintain NAD(H) nucleotides in the linear range of the calibration curve. Samples for NADPH and NADP^+^ estimation were not diluted as flies contain ∼10 times smaller NADP pool compared to NAD pool. Calibration curves were constructed for NAD^+^, NADH, NADP^+^ and NADPH nucleotides prepared in the dilution buffer containing 10 μM ascorbic acid. To each fifty μL of samples and standards in all-white 96-well plates, another 50 μL of the corresponding NAD/NADH-Glo or NADP/NADPH-Glo Detection Reagent (Promega cat. #: G9071 and G9081) were added, and luminescence was measured over 2 h using BioTek Cytation C10 Confocal Imaging Reader (Agilent Technologies). In our luminescence-based assays, we do not take end-time points. Instead, we monitor changes in luminescence over time and take a derivative of that change in the linear portion of the reaction progress curve to maximize the accuracy and reproducibility of our measurements (both samples and standards). In all figures, we report the total amount of nucleotides in pmol from 10 flies taking into account all dilutions during sample preparation.

### Immunoblotting

For immunoblot analysis, 5 adult flies were lysed in RIPA lysis buffer (Cell Signaling) with added phosphatase and protease inhibitors (Roche), using the bullet blender tissue homogenizer. The lysates were resolved on mini-Protean TGX gel (Bio-Rad) by electrophoresis followed by transfer on PVDF membrane (Immobilon P; Millipore), blocked in Tris-buffered saline Tween-20 buffer (Cell Signaling Technology) containing 2.5% dry milk and probed with the mentioned antibodies diluted in this buffer. For detecting *L*bNOX, anti-FLAG antibody 12C6c (DHSB) was used at 1:4000 dilution. For loading control, antibody against ²-Actin (13E5, Cell Signaling technology) was used at 1:1000 dilution.

### Metabolomics

Metabolic profiling was performed as previously described (69). Briefly, ten to twenty flies per sample (four biological replicates) were collected and intracellular metabolites were extracted using 80% (v/v) aqueous methanol. LC was performed on an Xbridge BEH amide HILIC column (Waters) with a Vanquish UHPLC system (Thermo Fisher). Solvent A was 95:5 water: acetonitrile with 20 mM ammonium acetate and 20 mM ammonium hydroxide at pH 9.4. Solvent B was acetonitrile. The gradient used for metabolite separation was 0 min, 90% B; 2 min, 90% B; 3 min, 75%; 7 min, 75% B; 8 min, 70% B, 9 min, 70% B; 10 min, 50% B; 12 min, 50% B; 13 min, 25% B; 14 min, 25% B; 16 min, 0% B, 21 min, 0% B; 21 min, 90% B; and 25 min, 90% B. MS analysis was performed on a Orbitrap Exploris 480 mass spectrometer (Thermo Fisher) by electrospray ionization with parameters as follow: scan mode, full MS; spray voltage, 3.6 kV (positive) and -3.2 kV (negative); capillary temperature, 320 °C; sheath gas, 40 arb; aux gas, 7 arb; resolution, 120,000 (full MS); scan *m*/*z* range 70 to 1,000. Web-based platform MetaboAnalyst was used to analyze and visualize metabolomics data.

### Sleep Measurements

*Drosophila* activity monitors (DAM) from Trikinetics Inc. (Waltham MA, USA) were used to monitor sleep and activity in flies. One-week or 3-weeks old flies (as mentioned in the text) were housed individually inside borosilicate glass tubes (65mm length, 5mm diameter) containing fly food at one end (coated with paraffin wax to avoid drying) and a cotton plug at the other end. Caution was taken while loading the 3-weeks old flies to not expose them to CO2 longer than 3 minutes while loading. Locomotor activity data were collected under LD (12 h light/12 h dark). Locomotor activity was recorded over a period of 4 days and sleep was analyzed using ShinyR-DAM (https://karolcichewicz.shinyapps.io/shinyr-dam/).

### NAD precursor supplementation

Flies were collected within 24-48 hrs from eclosion and allowed to mate for a day followed by sorting by sex under CO2 anesthesia. Post separations flies were placed on fly food containing specific NAD precursor. The flies were flipped onto fresh food vials every 2 days throughout the treatment. Following NAD precursors were used in the study: Nicotinic acid (NA) from Sigma-Aldrich catalog# N4126, N-acteyl cysteine (NAC) from Sigma-Aldrich catalog# A7250; sodium 3-hydroxybutyrate (BHB) from Sigma-Aldrich catalog# 54965; Nicotinamide riboside chloride (NR) from Selleckchem catalog# S2935 was used for NAD^+^ measurements; NR from Cayman chemical catalog# 36941 was used for lifespan analysis; Nicotinic Acid Mononucleotide (NMN) from Cayman chemical catalog# 32883.

## Acknowledgments

We thank Kent G. Golic for providing GMR-Gal4 UAS-CryAB^R120G^ stock. This work was supported by NIGMS R35 GM146869 (A.P.), NIA R01 AG082801 (A.P.), NIA R03 AG075651 (A.P.), R03 CA286521 (A.P.), NIA P30 AG024827 pilot grant (A.P.), Richard King Mellon Foundation award (A.P.), NAM Healthy Longevity Catalyst Award (A.P.), NIA R03AG067301 (V.C.), NIA R21AG085012 (V.C.), NIGMS R35GM142495 (V.C), and Longevity Impetus Grants (V.C.). This is manuscript number 1078 from The Scintillon Institute.

## Data and materials availability

All data are available within the Article and Supplementary Files, or available from the authors upon a reasonable request.

---

**Supplementary Figure S1:**
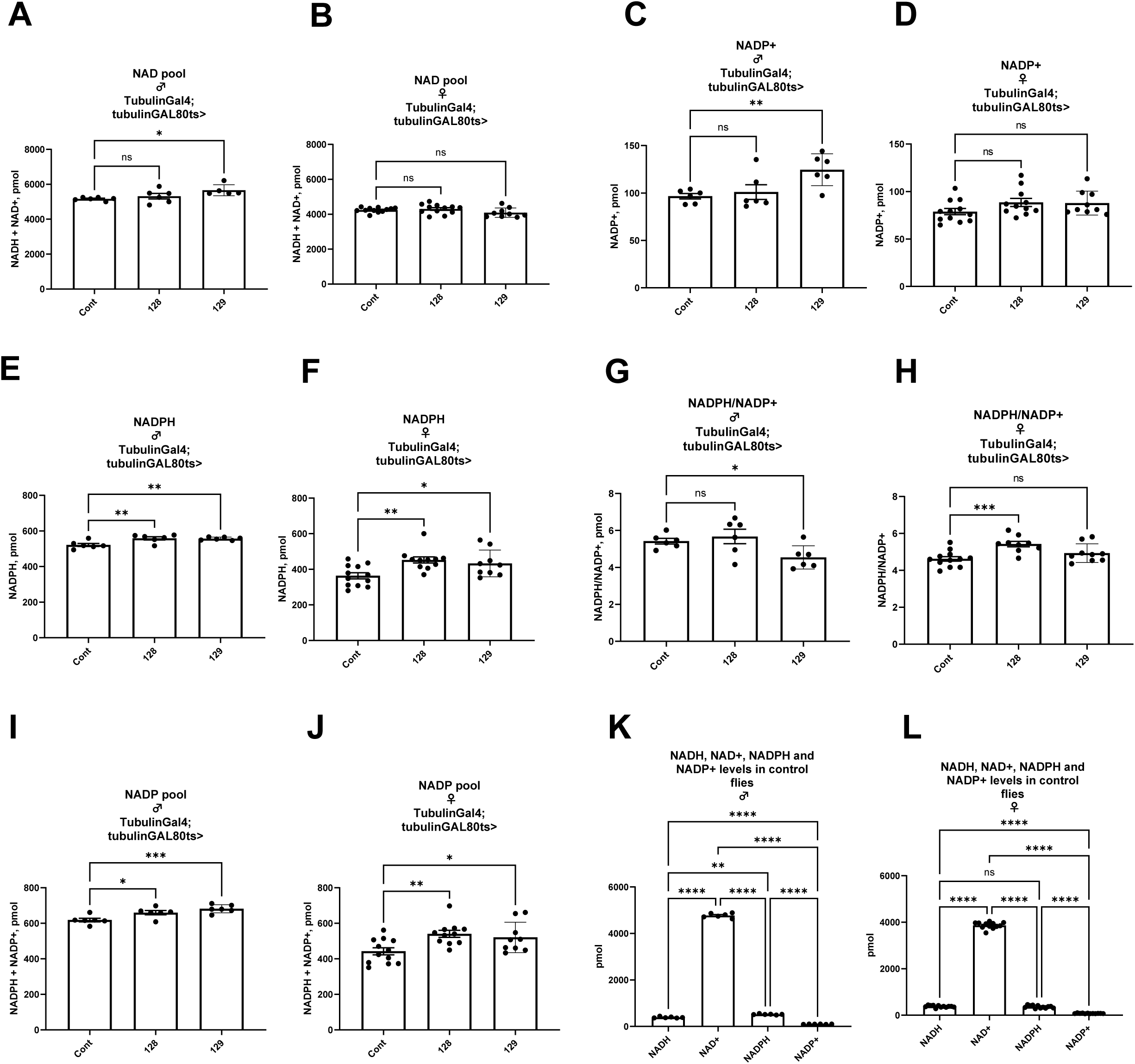
*Lb*NOX expression alters NAD(P)H/NAD(P)^+^ ratios in *Drosophila* in a sex-dependent manner. NAD pool (NAD^+^ + NADH) in males **(A)** and females (**B**) of *Drosophila* lines 128 and 129 expressing *Lb*NOX. Levels of NADP^+^ (**C**), NADPH (**E**), NADPH/NADP^+^ ratio (**G**) and NADP pool (NADP^+^ + NADPH) (**I**) in males of *Drosophila* lines 128 and 129 expressing *Lb*NOX. Levels of NADP^+^ (**D**), NADPH (**F**), NADPH/NADP^+^ ratio (**H**) and NADP pool (NADP^+^ + NADPH) (**J**) in females *Drosophila* lines 128 and 129 expressing *Lb*NOX. Levels of NADH, NAD^+^, NADPH and NADP^+^ shown for control males (**K**) and females (**L**) *Drosophila*. Values indicated in (A-L) are from 10 flies. The statistical significance indicated for (A-L) represents a one-way ANOVA followed by uncorrected Fishers Least Significant Difference (LSD) test. The error bars in (A-L) represent mean ± s.d.

**Supplementary Figure S2:**
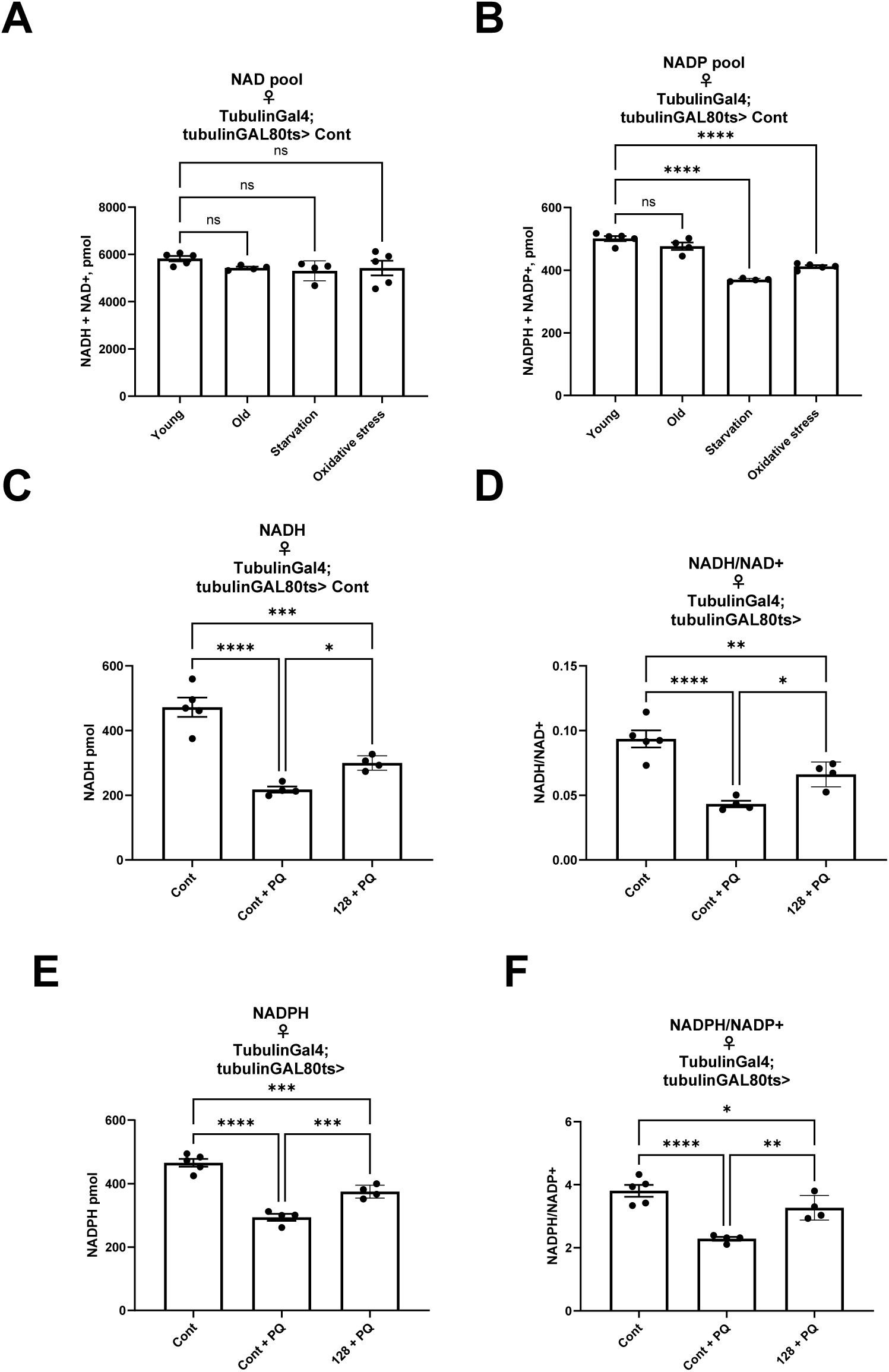
*Lb*NOX expression protects against detrimental effects of oxidative stress. NAD pool (NAD^+^ + NADH) (**A**) and NADP pool (NADP^+^ + NADPH) (**B**) in female flies that were 1 week old(young), 3 weeks old (old), subjected to starvation or oxidative stress for 16 hrs. Levels of NADH (**C**), NADH/NAD^+^ ratio (**D**), NADPH (**E**) and NADPH/NADP^+^ ratio (**F**) in female controls and female *Lb*NOX line 128 flies fed with food containing 10mM of paraquat (PQ) for 16hrs. Values indicated in (A-F) are from 10 flies. The statistical significance indicated for (A-F) represents a one-way ANOVA followed by uncorrected Fishers Least Significant Difference (LSD) test. The error bars in (A-F) represent mean ± s.d.

**Supplementary Figure S3:**
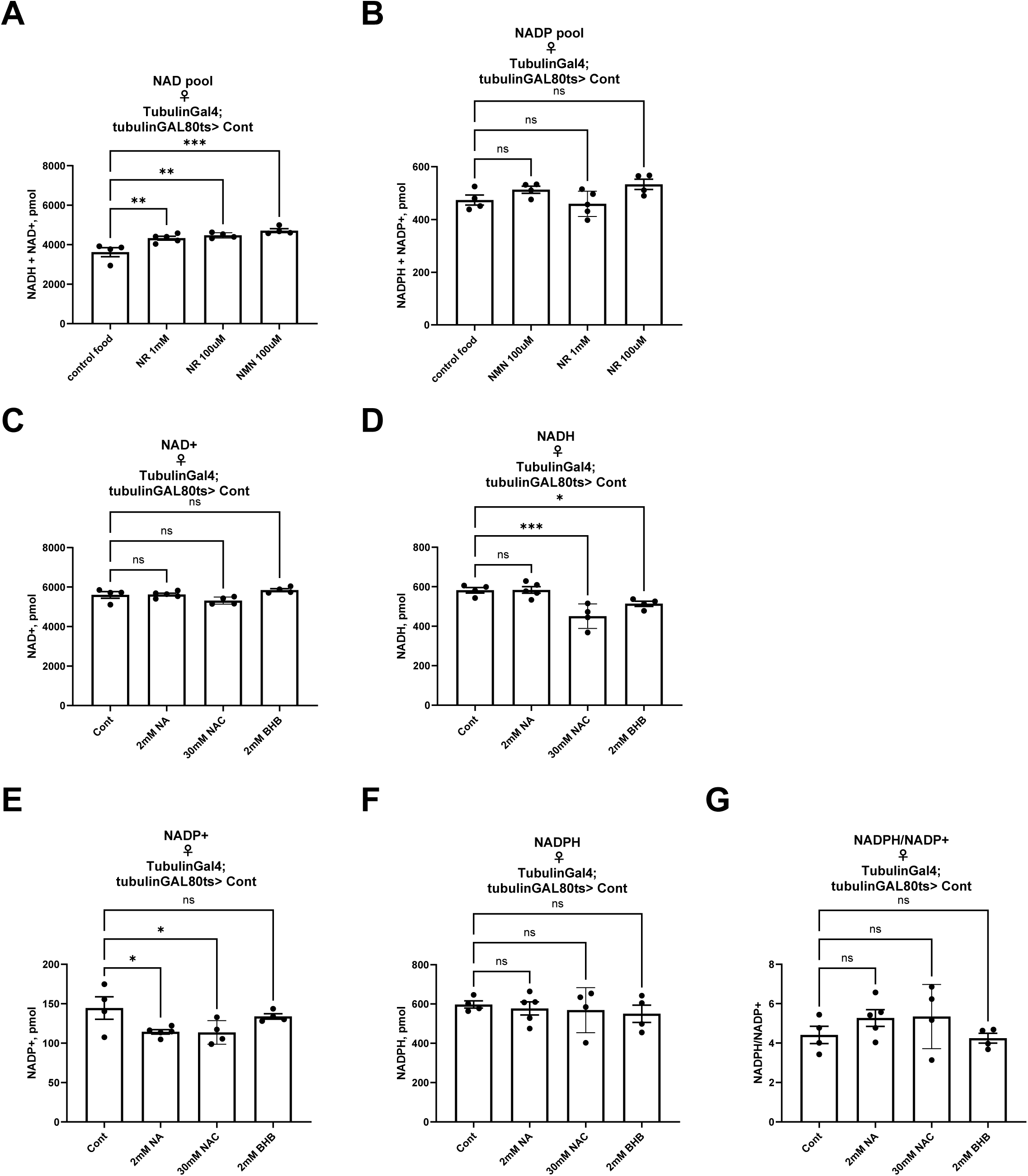
NAD precursor supplementations do not further extend lifespan of *Drosophila* expressing *Lb*NOX ubiquitously. NAD pool (NAD^+^ + NADH) (**A**) and NADP pool (NADP^+^ + NADPH) (**B**) in female control flies feed with food containing 1mM NR, 100μM NR or 100μM NMN for 7 days. Levels of NAD^+^ (**C**), NADH (**D**), NADP^+^ (**E**), NADPH (**F**) and NADPH/NADP^+^ ratio (**G**) in control female flies fed with food with 2mM NA, 30mM NAC or 2mM BHB compared to flies fed on control food. Values indicated in (A-G) are from 10 flies. The statistical significance indicated for (A-G) represents a one-way ANOVA followed by uncorrected Fishers Least Significant Difference (LSD) test. The error bars in (A-G) represent mean ± s.d.

**Supplementary Figure S4:**
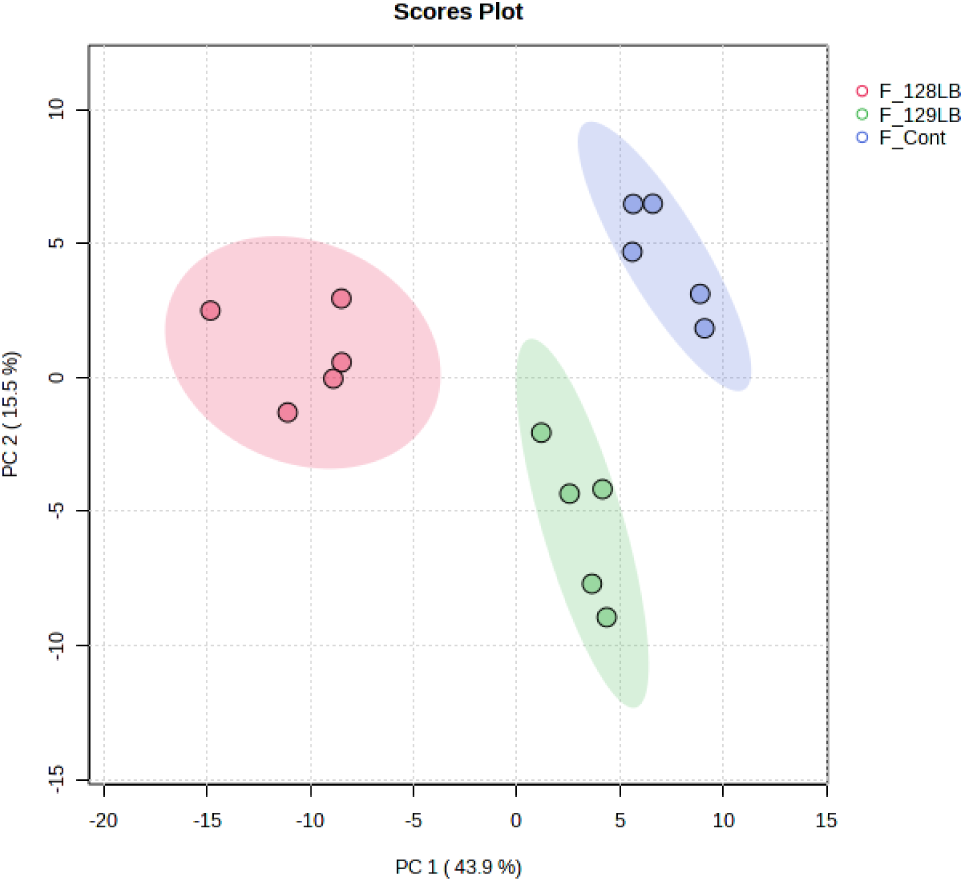
*Lb*NOX expression reprograms *Drosophila* metabolism with a strong impact on *de novo* NAD biosynthesis. Principal component analysis of metabolic profiling of female flies expressing *Lb*NOX lines 128 and 129.

**Supplementary Figure S5:**
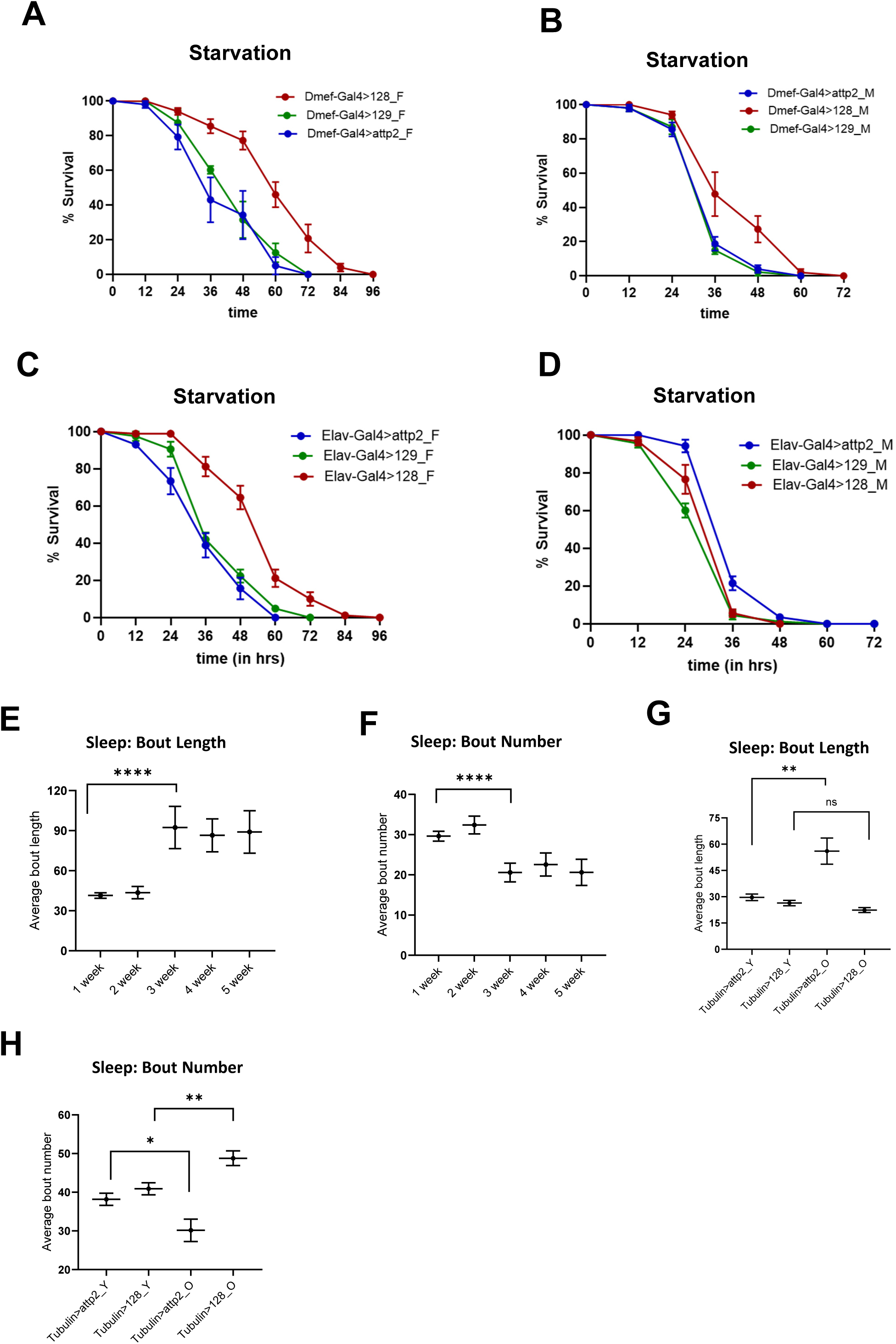
Tissue-specific *Lb*NOX expression efficiently protects against oxidative stress and rejuvenates sleep profiles in aged flies back to a youthful state. Survival of female (**A**) and male (**B**) control flies and flies with muscle-specific *Lb*NOX expression under starvation. Survival of female (**C**) and male (**D**) control flies and flies with neuron-specific *Lb*NOX expression under starvation. Sleep bout length **(E)** and bout number **(F)** in female control flies of mentioned ages (1 week through 5 weeks). Sleep bout length **(E)** and bout number **(F)** in control flies or flies with ubiquitous expression of LbNOX #128 at 1-week(marked as Y) and 3-weeks (marked as O) of age. The error bars in (A-D) represent mean ± s.e.m.

## Supplementary Tables

**Supplementary Table S1: Summary of NAD(P)(H) metabolites in flies expressing *Lb*NOX 128 and 129 lines and their associated phenotypes.**

Summary of NAD(P)(H) metabolites and their ratios in flies ubiquitously expressing *Lb*NOX #128 and #129 of both sexes.

**Supplementary Table S2: Significantly altered metabolites in flies expressing *Lb*NOX 128 and 129 lines.**

Significantly altered metabolites in flies ubiquitously expressing *Lb*NOX #128 and #129 of both sexes.

**Supplementary Table S3: Metabolite enrichment analysis on significantly altered metabolites in flies expressing *Lb*NOX 128 and 129 lines.**

Metabolite enrichment analysis on significantly altered metabolites in flies ubiquitously expressing *Lb*NOX #128 and #129 of both sexes.

**Supplementary Table S4: Key resources table.**

Key resources table listing all *Drosophila* lines, kits, antibodies, chemicals, and other consumables.

## References

1. A. A. Parkhitko, E. Filine, S. E. Mohr, A. Moskalev, N. Perrimon, Targeting metabolic pathways for extension of lifespan and healthspan across multiple species. Ageing Res Rev 64, 101188 (2020).

2. E. Katsyuba, M. Romani, D. Hofer, J. Auwerx, NAD(+) homeostasis in health and disease. Nat Metab 2, 9–31 (2020).

3. S. Lautrup, D. A. Sinclair, M. P. Mattson, E. F. Fang, NAD(+) in Brain Aging and Neurodegenerative Disorders. Cell Metab 30, 630–655 (2019).

4. A. J. Covarrubias, R. Perrone, A. Grozio, E. Verdin, NAD(+) metabolism and its roles in cellular processes during ageing. Nat Rev Mol Cell Biol 22, 119–141 (2021).

5. W. Xiao, R. S. Wang, D. E. Handy, J. Loscalzo, NAD(H) and NADP(H) Redox Couples and Cellular Energy Metabolism. Antioxid Redox Signal 28, 251–272 (2018).

6. C. C. S. Chini, J. D. Zeidler, S. Kashyap, G. Warner, E. N. Chini, Evolving concepts in NAD(+) metabolism. Cell Metab 33, 1076–1087 (2021).

7. A. Peluso, M. V. Damgaard, M. A. S. Mori, J. T. Treebak, Age-Dependent Decline of NAD(+)-Universal Truth or Confounded Consensus? Nutrients 14 (2021).

8. L. Rajman, K. Chwalek, D. A. Sinclair, Therapeutic Potential of NAD-Boosting Molecules: The In Vivo Evidence. Cell Metab 27, 529–547 (2018).

9. M. V. Damgaard, J. T. Treebak, What is really known about the effects of nicotinamide riboside supplementation in humans. Sci Adv 9, eadi4862 (2023).

10. D. V. Titov et al., Complementation of mitochondrial electron transport chain by manipulation of the NAD+/NADH ratio. Science 352, 231–235 (2016).

11. W. Xiao, J. Loscalzo, Metabolic Responses to Reductive Stress. Antioxid Redox Signal 32, 1330–1347 (2020).

12. E. C. Lien et al., Effects of Aging on Glucose and Lipid Metabolism in Mice. Aging Cell 10.1111/acel.14462, e14462 (2024).

13. S. E. McGuire, P. T. Le, A. J. Osborn, K. Matsumoto, R. L. Davis, Spatiotemporal rescue of memory dysfunction in Drosophila. Science 302, 1765–1768 (2003).

14. M. Mladenov et al., Oxidative Stress, Reductive Stress and Antioxidants in Vascular Pathogenesis and Aging. Antioxidants (Basel) 12 (2023).

15. A. G. Manford et al., A Cellular Mechanism to Detect and Alleviate Reductive Stress. Cell 183, 46–61 e21 (2020).

16. H. B. Xie et al., The NADPH metabolic network regulates human alphaB-crystallin cardiomyopathy and reductive stress in Drosophila melanogaster. PLoS Genet 9, e1003544 (2013).

17. P. Vicart et al., A missense mutation in the alphaB-crystallin chaperone gene causes a desmin-related myopathy. Nat Genet 20, 92–95 (1998).

18. C. R. Reczek et al., A CRISPR screen identifies a pathway required for paraquat-induced cell death. Nat Chem Biol 13, 1274–1279 (2017).

19. G. Roman, K. Endo, L. Zong, R. L. Davis, P[Switch], a system for spatial and temporal control of gene expression in Drosophila melanogaster. Proceedings of the National Academy of Sciences of the United States of America 98, 12602–12607 (2001).

20. T. Osterwalder, K. S. Yoon, B. H. White, H. Keshishian, A conditional tissue-specific transgene expression system using inducible GAL4. Proceedings of the National Academy of Sciences of the United States of America 98, 12596–12601 (2001).

21. E. F. Fang et al., NAD(+) augmentation restores mitophagy and limits accelerated aging in Werner syndrome. Nat Commun 10, 5284 (2019).

22. K. Richardson, R. Wessells, A novel panel of Drosophila TAFAZZIN mutants in distinct genetic backgrounds as a resource for therapeutic testing. PLoS One 18, e0286380 (2023).

23. N. C. Yang, Y. H. Cho, I. Lee, The Lifespan Extension Ability of Nicotinic Acid Depends on Whether the Intracellular NAD(+) Level Is Lower than the Sirtuin-Saturating Concentrations. Int J Mol Sci 21 (2019).

24. V. Balan et al., Life span extension and neuronal cell protection by Drosophila nicotinamidase. J Biol Chem 283, 27810–27819 (2008).

25. F. Cavaleri, E. Bashar, Potential Synergies of beta-Hydroxybutyrate and Butyrate on the Modulation of Metabolism, Inflammation, Cognition, and General Health. J Nutr Metab 2018, 7195760 (2018).

26. A. Fiore, P. J. Murray, Tryptophan and indole metabolism in immune regulation. Curr Opin Immunol 70, 7–14 (2021).

27. C. Xue et al., Tryptophan metabolism in health and disease. Cell Metab 35, 1304–1326 (2023).

28. M. De Giovanni, H. Chen, X. Li, J. G. Cyster, GPR35 and mediators from platelets and mast cells in neutrophil migration and inflammation. Immunol Rev 317, 187–202 (2023).

29. M. Kaihara, J. M. Price, The metabolism of quinaldic acid, kynurenic acid, and xanthurenic acid in the rabbit. J Biol Chem 237, 1727–1729 (1962).

30. R. Castro-Portuguez, G. L. Sutphin, Kynurenine pathway, NAD(+) synthesis, and mitochondrial function: Targeting tryptophan metabolism to promote longevity and healthspan. Exp Gerontol 132, 110841 (2020).

31. V. Mariano et al., SREBP modulates the NADP(+)/NADPH cycle to control night sleep in Drosophila. Nat Commun 14, 763 (2023).

32. A. Kempf, S. M. Song, C. B. Talbot, G. Miesenbock, A potassium channel beta-subunit couples mitochondrial electron transport to sleep. Nature 568, 230–234 (2019).

33. P. N. Bushana, M. A. Schmidt, M. J. Rempe, B. A. Sorg, J. P. Wisor, Chronic dietary supplementation with nicotinamide riboside reduces sleep need in the laboratory mouse. Sleep Adv 4, zpad044 (2023).

34. K. M. Ramsey et al., Circadian clock feedback cycle through NAMPT-mediated NAD+ biosynthesis. Science 324, 651–654 (2009).

35. M. Morifuji, S. Higashi, S. Ebihara, M. Nagata, Ingestion of beta-nicotinamide mononucleotide increased blood NAD levels, maintained walking speed, and improved sleep quality in older adults in a double-blind randomized, placebo-controlled study. Geroscience 46, 4671–4688 (2024).

36. B. Cuenoud et al., Effect of circadian rhythm on NAD and other metabolites in human brain. Front Physiol 14, 1285776 (2023).

37. T. S. Andreani, T. Q. Itoh, E. Yildirim, D. S. Hwangbo, R. Allada, Genetics of Circadian Rhythms. Sleep Med Clin 10, 413–421 (2015).

38. C. L. Partch, C. B. Green, J. S. Takahashi, Molecular architecture of the mammalian circadian clock. Trends Cell Biol 24, 90–99 (2014).

39. A. Crocker, A. Sehgal, Genetic analysis of sleep. Genes Dev 24, 1220–1235 (2010).

40. E. J. Beckwith, A. S. French, Sleep in Drosophila and Its Context. Front Physiol 10, 1167 (2019).

41. A. Sanz et al., Expression of the yeast NADH dehydrogenase Ndi1 in Drosophila confers increased lifespan independently of dietary restriction. Proceedings of the National Academy of Sciences of the United States of America 107, 9105–9110 (2010).

42. S. Bahadorani et al., Neuronal expression of a single-subunit yeast NADH-ubiquinone oxidoreductase (Ndi1) extends Drosophila lifespan. Aging Cell 9, 191–202 (2010).

43. K. K. Kemppainen et al., Expression of alternative oxidase in Drosophila ameliorates diverse phenotypes due to cytochrome oxidase deficiency. Hum Mol Genet 23, 2078–2093 (2014).

44. D. J. Fernandez-Ayala et al., Expression of the Ciona intestinalis alternative oxidase (AOX) in Drosophila complements defects in mitochondrial oxidative phosphorylation. Cell Metab 9, 449–460 (2009).

45. A. A. Parkhitko et al., A genetic model of methionine restriction extends Drosophila health-and lifespan. Proceedings of the National Academy of Sciences of the United States of America 118 (2021).

46. V. Cracan, D. V. Titov, H. Shen, Z. Grabarek, V. K. Mootha, A genetically encoded tool for manipulation of NADP(+)/NADPH in living cells. Nat Chem Biol 13, 1088–1095 (2017).

47. X. Pan et al., A genetically encoded tool to increase cellular NADH/NAD(+) ratio in living cells. Nat Chem Biol 20, 594–604 (2024).

48. R. Yang et al., Identification of purine biosynthesis as an NADH-sensing pathway to mediate energy stress. Nat Commun 13, 7031 (2022).

49. A. A. Parkhitko et al., Cross-species identification of PIP5K1-, splicing-and ubiquitin-related pathways as potential targets for RB1-deficient cells. PLoS Genet 17, e1009354 (2021).

50. S. Yadav et al., Unique tau-and synuclein-dependent metabolic reprogramming in neurons distinct from normal aging. Aging Cell 23, e14277 (2024).

51. A. A. Parkhitko, E. Filine, M. Tatar, Combinatorial interventions in aging. Nat Aging 3, 1187–1200 (2023).

52. A. H. Brand, N. Perrimon, Targeted gene expression as a means of altering cell fates and generating dominant phenotypes. Development 118, 401–415 (1993).

53. M. R. McReynolds et al., NAD(+) flux is maintained in aged mice despite lower tissue concentrations. Cell Syst 12, 1160–1172 e1164 (2021).

54. S. Rimal et al., Reverse electron transfer is activated during aging and contributes to aging and age-related disease. EMBO Rep 24, e55548 (2023).

55. R. P. Goodman, S. E. Calvo, V. K. Mootha, Spatiotemporal compartmentalization of hepatic NADH and NADPH metabolism. J Biol Chem 293, 7508–7516 (2018).

56. H. Y. Ho, Y. T. Lin, G. Lin, P. R. Wu, M. L. Cheng, Nicotinamide nucleotide transhydrogenase (NNT) deficiency dysregulates mitochondrial retrograde signaling and impedes proliferation. Redox Biol 12, 916–928 (2017).

57. L. Del Prado et al., Compensatory activity of the PC-ME1 metabolic axis underlies differential sensitivity to mitochondrial complex I inhibition. Nat Commun 15, 8682 (2024).

58. E. Balsa et al., Defective NADPH production in mitochondrial disease complex I causes inflammation and cell death. Nat Commun 11, 2714 (2020).

59. S. Igelmann et al., A hydride transfer complex reprograms NAD metabolism and bypasses senescence. Mol Cell 81, 3848–3865 e3819 (2021).

60. P. C. Bradshaw, Cytoplasmic and Mitochondrial NADPH-Coupled Redox Systems in the Regulation of Aging. Nutrients 11 (2019).

61. O. M. Bubu et al., Sleep, Cognitive impairment, and Alzheimer’s disease: A Systematic Review and Meta-Analysis. Sleep 40 (2017).

62. A. Iranzo, Sleep in Neurodegenerative Diseases. Sleep Med Clin 11, 1–18 (2016).

63. J. E. Carroll, A. A. Prather, Sleep and Biological Aging: A Short Review. Curr Opin Endocr Metab Res 18, 159–164 (2021).

64. S. Dissel, Drosophila as a Model to Study the Relationship Between Sleep, Plasticity, and Memory. Front Physiol 11, 533 (2020).

65. J. M. Lane et al., Biological and clinical insights from genetics of insomnia symptoms. Nat Genet 51, 387–393 (2019).

66. L. Schwarzmann, R. U. Pliquett, A. Simm, B. Bartling, Sex-related differences in human plasma NAD+/NADH levels depend on age. Biosci Rep 41 (2021).

67. V. van der Velpen et al., Sex-specific alterations in NAD+ metabolism in 3xTg Alzheimer’s disease mouse brain assessed by quantitative targeted LC-MS. J Neurochem 159, 378–388 (2021).

68. N. S. Rajasekaran et al., Human alpha B-crystallin mutation causes oxido-reductive stress and protein aggregation cardiomyopathy in mice. Cell 130, 427–439 (2007).

69. A. A. Parkhitko et al., Downregulation of the tyrosine degradation pathway extends Drosophila lifespan. Elife 9 (2020).

