## Supplementary Table S1 for "Tissue-specific modulation of NADH consumption as an anti-aging intervention in Drosophila"

| Parameter | UAS>128 | | UAS>129 | |
|---|---|---|---|---|
| | Female | Male | Female | Male |
| **Alteration in NAD metabolites levels** | | | | |
| NAD+ | -- | -- | -- | Inc |
| NADP+ | -- | -- | -- | Inc * |
| NADH | -- | Dec | -- | Dec |
| NADPH | Inc ** | Inc ** | Inc * | Inc ** |
| **Total pool of metabolite** | | | | |
| NAD pool | -- | -- | -- | Inc * |
| NADP pool | Inc ** | Inc * | Inc * | Inc *** |
| **Stress tolerance** | | | | |
| Starvation stress | R | -- | R | -- |
| Oxidative stress | R | R | R | -- |
| **Impact on lifespan when expressed with** | | | | |
| Da-GS-GAL4 | E | -- | E | -- |
| Actin-GS-GAL4 | E | -- | -- | -- |
| **Stress tolerance upon tissue specific expression** | | | | |
| Muscle-specific (Dmef-GAL4) | | | | |
| Starvation stress | R | R | -- | -- |
| Oxidative stress | R | -- | R | -- |
| Neuron specific (Elav-GAL4) | | | | |
| Starvation stress | R | -- | S | S |
| Oxidative stress | R | S | R | S |

**Table 1**. Summary of changes induced by expression of LbNOX using UAS-GAL4 system. Abbreviations used are Inc: increase, Dec: decrease, E: extension, R: resistant, S: sensitive.
