## Supplementary Table S2 for "Tissue-specific modulation of NADH consumption as an anti-aging intervention in Drosophila"

**Table 2**: Significantly altered metabolites in flies expressing LbNOX 128 and 129 lines.

| 128 | 129 | 128\|129 shared |
|---|---|---|
| glutathione | 5-hydroxyisourate | Indole-3-acetic acid |
| hydroxypyruvate | Indole-3-acetic acid | Xanthurenic acid |
| N-Methyl-L-lysine | Xanthurenic acid | Tryptamine |
| 3-Hydroxypropionic Acid | Tryptamine | Indole-3-carboxylic acid |
| carbamoyl phosphate | Indole-3-carboxylic acid | tryptophan |
| lactate | tryptophan | 5-hydroxyindole-3-acetic acid |
| cyclic AMP | 5-hydroxyindole-3-acetic acid | Quinaldic acid |
| hydroxyphenylpyruvate | Quinaldic acid | kynurenine |
| sorbitol | kynurenine | Kynurenic acid |
| allantoin | Kynurenic acid | Indoleacetic acid |
| Methyl phosphoethanolamine | Indoleacetic acid | Hippuric acid |
| 3_4-Dihydroxyhydrocinnamic acid | Hippuric acid | Adonitol |
| lysine | Adonitol | Malonic acid |
| Pantetheine | Malonic acid | GDP-glucose |
| Deoxyinosine monophosphate | GDP-glucose | ribose-5-phosphate |
| adenine | ribose-5-phosphate | 1-Methyladenosine |
| orotate | 1-Methyladenosine | CDP-ethanolamine |
| 3-Hydroxyphenylacetic acid | CDP-ethanolamine | xanthine |
| N-acetyl-glutamine | xanthine | |
| C20:0 | | |
| tyrosine | | |
| Methionine sulfoxide | | |
| Saccharopine | | |
| D-Galactonic acid | | |
| C18:1 | | |
| uridine | | |
| 4-Acetamidobutyric acid | | |
| D-phospho-L-serine | | |
| 3-phosphoglycerate | | |
| C14:0 | | |
| a-ketoglutarate | | |
| Glycerylphosphorylethanolamine | | |
| glycerate | | |
| CMP | | |
| LysoPE(18:1) | | |
| glucosamine-1-phosphate | | |
| Heptanoic acid | | |
| C16:1 | | |
| N-Acetylasparagine | | |
| alpha-Ketoglutaramate | | |
| phosphoenolpyruvate | | |
| 6-Keto-decanoylcarnitine | | |
| VaL-Gly | | |
| citrate/isocitrate | | |
| N-Acetylvaline | | |
| IMP | | |
| 2-Amino-3-phosphonopropionic acid | | |
| NADP | | |
| GMP | | |
| N-Formyl-L-aspartate | | |
| Acetylglycine | | |
| pyruvate | | |
