## Supplementary Table S3 for "Tissue-specific modulation of NADH consumption as an anti-aging intervention in Drosophila"

**Table 3**: Metabolite enrichment analysis on significantly altered metabolites in flies expressing LbNOX 128 and 129 lines.

| Term | total | expected | hits | Raw p | X.LOG10.p. | Holm p | FDR | |
|---|---|---|---|---|---|---|---|---|
| Citrate cycle (TCA cycle) | 20 | 0.234 | 4 | 5.66E-05 | 4.24718357 | 0.00453 | 0.00453 | Up |
| Alanine, aspartate and glutamate metabolism | 28 | 0.327 | 4 | 0.000226 | 3.64589156 | 0.0178 | 0.00902 | Up |
| Glyoxylate and dicarboxylate metabolism | 31 | 0.363 | 3 | 0.00492 | 2.3080349 | 0.384 | 0.118 | Up |
| Glycine, serine and threonine metabolism | 33 | 0.386 | 3 | 0.00589 | 2.22988471 | 0.453 | 0.118 | Up |
| Tryptophan metabolism | 41 | 0.506 | 5 | 0.0000925 | 4.03385827 | 0.0074 | 0.0074 | Down |
| Phenylalanine, tyrosine and tryptophan biosynthe | 4 | 0.0494 | 2 | 0.000854 | 3.06854213 | 0.0675 | 0.0342 | Down |
| Ubiquinone and other terpenoid-quinone biosynt | 18 | 0.222 | 2 | 0.0196 | 1.70774393 | 1 | 0.524 | Down |
| Lysine degradation | 30 | 0.37 | 2 | 0.0512 | 1.29073004 | 1 | 0.828 | Down |
