## Supplementary Table S4 for "Tissue-specific modulation of NADH consumption as an anti-aging intervention in Drosophila"

### Table 4. Key resources table

| Reagent type | Designation | Source or reference | Identifiers | Additional Information |
|---|---|---|---|---|
| Genetic reagent (*D.melanogaster*) | tubulinGAL4; tubulinGal80$^{ts}$ | Perrimon's lab | | Used for whole body expression |
| Genetic reagent (*D.melanogaster*) | Elav-GAL4; tubulinGal80$^{ts}$ | Perrimon's lab | | Used for neuronal/head expression |
| Genetic reagent (*D.melanogaster*) | Dmef-GAL4; tubulinGal80$^{ts}$ | Perrimon's lab | | Used for muscle-specific expression |
| Genetic reagent (*D.melanogaster*) | GMR-GAL4 | Perrimon's lab | | Used for expression in eyes |
| Genetic reagent (*D.melanogaster*) | GMR>cryAB | Gift from Dr. Kent G. Golic | | Model for reductive stress |
| Genetic reagent (*D.melanogaster*) | *attp2* | Bloomington *Drosophila* stock center (BDSC) | 36303 | Used as a control |
| Genetic reagent (*D.melanogaster*) | *Actin-GS-Gal4* | | | GeneSwitch line |
| Genetic reagent (*D.melanogaster*) | *Da-GS-Gal4* | | | GeneSwitch line |
| Plasmid construct | pWALIUM10-ROE | | | Used for cloning *Lb*NOX |
| Kits | NAD/NADH-Glo or NADP/NADPH-Glo Detection Reagent | Promega | G9071 and G9081 | Used for $NAD^+$, NADH, $NADP^+$ and NADPH estimation |
| anti-FLAG antibody | | DHSB | 12C6c | For *Lb*NOX detection in WB |
| Anti-β-Actin antibody | | CST | 13E5 | For actin detection in WB |
| Chemical | Nicotinic acid | Sigma-Aldrich | N4126 | For supplementation |
| Chemical | N-acteyl cysteine | Sigma-Aldrich | A7250 | For supplementation |
| Chemical | Sodium 3-hydroxybutyrate | Sigma-Aldrich | 54965 | For supplementation |
| Chemical | Nicotinamide riboside chloride | Selleckchem | S2935 | For supplementation |
| Chemical | Nicotinamide riboside chloride | Cayman | 36941 | For lifespan analysis |
| Chemical | Nicotinic Acid Mononucleotide | Cayman | 32883 | For supplementation |
